# Laws for Glia Organization Conserved Across Mammals

**DOI:** 10.1101/449421

**Authors:** Antonio Pinto-Duarte, Katharine Bogue, Terrence J. Sejnowski, Shyam Srinivasan

**Affiliations:** Computational Neurobiology Laboratory, Salk Institute for Biological Studies; La Jolla, CA 92037; Molecular Neurobiology Laboratory, Salk Institute for Biological Studies; La Jolla, CA 92037; Institute for Neural Computation, University of California, San Diego; 9500 Gilman Drive, CA 92093; Division of Biological Sciences, University of California, San Diego; CA 92093

**Keywords:** Brain size, glia density, scaling, brain organization, glia-neuron ratio

## Abstract

The organizational principles of glia remain largely unknown despite their vital role in nervous system function. Previous work has shown that the number of glia per unit volume of neocortex is constant across mammalian species. We hypothesize that the conservation of glia volume density within brain regions might be a governing principle of organization across species. To test this hypothesis, we used stereology, light microscopy, and data available in the literature to examine five brain regions: the cerebral cortex and four brain regions that differ from the cerebral cortex and each other - the anterior piriform cortex, the posterior piriform cortex, the entorhinal cortex, and the cerebellum. We discovered two orderly relationships: First, glia volume density within a brain region was constant across species, including humans, although it significantly differed between regions, suggesting that glia density might constitute a region-specific marker. Second, the ratio of glia to neuron increased with brain volume according to a ¼ power law in the primate frontal cortex and the neocortex, the mammalian paleocortex, and the cerebellum. These relationships show that the development of glia and neurons are coupled, and suggest that what a neural circuit computes depends as much on its glial components as on its neurons.

**Main Points:**

- The volume density of glia (i.e., number of glia per unit volume) within a brain region is conserved across mammalian species including humans.
- The ratio of glia to neuron increases with bigger brains.
- The volume density of glia is significantly different across functionally and architecturally different brain regions and could function as a region-specific marker.
- Glia obey scaling constraints that are different from scaling constraints for neurons.

## Introduction

Numerous studies have examined how neural components in different brain regions scale with brain size (Finlay and Darlington, 1995a; Zhang and Sejnowski, 2000; Stevens, 2001; Cuntz et al., 2012; Buzsáki et al., 2013), a strategy which has contributed valuable insights into circuit function by linking architecture, information coding, and function (Stevens, 2001; Srinivasan et al., 2015; Srinivasan and Stevens, 2019). Most of those reports, however, have ignored glia as a whole, and current knowledge of how their absolute number (or their number relative to neurons) changes within a region, is limited to a few species and brain regions, e.g., the neocortex (Carlo and Stevens, 2013). Together with qualitative studies (Allen and Eroglu, 2017; Allen and Lyons, 2018; Kuhn et al., 2019; Santello et al., 2019) implicating glia in synaptic transmission and plasticity, neuronal regulation, immune responses, and brain development, quantitative studies are essential not only for understanding glia’s circuit-specific roles, but also for obtaining a comprehensive view on how information is processed in neural circuits.

Two quantitative metrics within a brain region -- glia densities and glia-to-neuron ratios (GNR) -- can inform our understanding of the roles played by glia in maintaining the health and information processing abilities of neuronal circuits (Sherwood et al., 2006; Herculano-Houzel, 2014). Neuronal circuits can be differentiated based on their architecture, connectivity, and the computations they perform, e.g., visual vs. olfactory circuits. Each circuit has diverse glial cell types, distinguishable on the basis of their morphology, function, and role (Allen and Lyons, 2018). Examples include astrocytes – which are implicated in critical functions, such as metabolism, synchronization of neuronal networks, and fine-tuning of synaptic transmission and plasticity (Mu et al., 2019; Pinto-Duarte et al., 2019; Santello et al., 2019); microglia, which play a key role in local immune responses through phagocytosis; and oligodendrocytes, which enhance neuronal communication by providing axon myelination (Allen and Lyons, 2018). As functional aspects, such as the amount of synchronization, synaptic plasticity, and information processed, differ between circuits, it is likely that the specific demands for local glial populations also differ. The quantification of glia densities reflect these differences.

Several studies have shown that GNR estimations can provide important insights into neuronal function (Sherwood et al., 2006; Herculano-Houzel, 2014). GNR estimates support the notion that more glia are required to accommodate the needs of more complex neuronal processes and computations in bigger brains (Sherwood et al., 2006). While existing evidence of increased GNR in the prefrontal cortex of bigger brains seems to support such a possibility (Sherwood et al., 2006), it is unknown whether GNR increases occur in architecturally different neural circuits. Concurrently, GNR estimations could deepen insights from previous findings of glia-neuron interactions such as neuronal regulation of astrocyte maturation and calcium signaling (Allen and Eroglu, 2017) or oligodendrocyte number and their activity (Barres and Raff, 1999; Simons and Trajkovic, 2006). For these reasons, a quantitative characterization of glia density and GNRs of different brain circuits is needed.

Therefore, we chose four regions that will help in achieving a fuller representation of the diversity in brain circuits. Circuits can be distinguished on the basis of their (i) connectivity -- topographic (there is a map or connections are predictable, e.g., the visual circuit where nearby neurons in the retina connect to nearby neurons in V1) or distributed (connections are unpredictable and are made without a spatial preference, i.e., there is no map, e.g., connections from OB to PCx), (ii) lamination – the number of layers, (iii) function – do they process a single sensory or multimodal input, and (iv) size of the circuit in terms of neuron numbers. Based on these factors, we examined the anterior piriform cortex or APCx, posterior piriform cortex or PPCx, the entorhinal cortex or ECx, and the cerebellum; regions that are significantly distinct from the well characterized neocortex, and each other. For instance, in the piriform cortex, afferent inputs from the olfactory bulb synapse onto, and activate, neurons distributed throughout the piriform, without any spatial preference (i.e., unpredictable), in contrast to regions in the neocortex like V1 that get topographic input from the retina. The regions are distinct in other ways, too. The cerebellum is anatomically separated from the cerebrum and contains 70% of the neurons in the human brain (Friede, 1963; Andersen et al., 1992, 2003)and a unique type of “specialized” radial astrocytes: the Bergmann glia. The anterior and posterior piriform cortices (Fig. 1a) are both trilaminar structures, comprising a sparsely populated layer 1, a densely packed layer 2, and an intermediately dense layer 3 (Neville and Haberly, 2004). Finally, the entorhinal cortex (ECx) is functionally distinct, as it is a multimodal circuit mixing input from topographic and distributed circuits and uses sensory information towards higher cognitive tasks such as spatial navigation (Fyhn et al., 2004; Witter et al., 2017).

**Figure 1:**
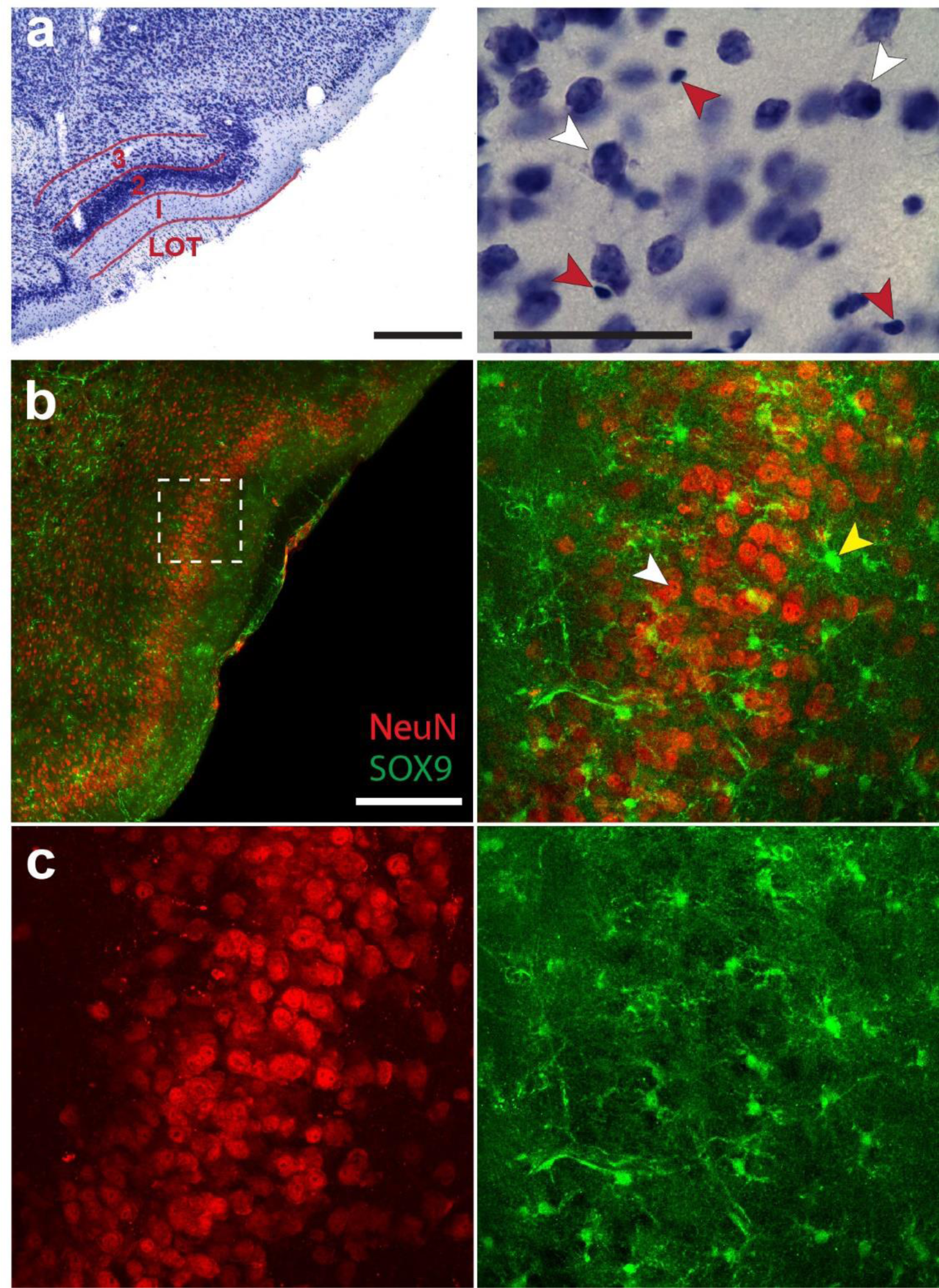
Identification of neurons, glia and astroglia in histological sections of the APCx. (a, left panel) Nissl-stained coronal section in a mouse showing the three-layered piriform cortex. Note that layer 2 is readily identifiable because of the extremely high density of neurons; layer 1 is cell-scarce, and layer 3 has intermediate cell density. Scale bar: 500 µm. (a, right panel) Magnified image of (a, 2x magnification) at a 100x magnification showing neurons and glia in Layer 3. Neurons (white arrowheads) are distinguishable from glia (red arrowheads) by their larger size and morphology (glia appear smaller and are more punctate, see methods for more detailed descriptions). (b, left panel) Example of a section of the APCx of eGFP-SOX9 mice co-labeled with the neuron marker, NeuN. Astrocytes and neurons are false-colored in green and red, respectively. The density of NeuN labelled cells clearly distinguishes Layer 2 from the other layers. Scale bar: 200 µm. Inset dotted line square (edge length: 200 µm) is shown on the right panel with arrow heads marking neurons and astrocytes (c) The left and right panels show the same image as (b, right panel) with only neurons (marked in red, left) or glia (marked in green, right).

We found two principles underlying glia organization. First, glia density is constant within the same circuit, independent of species or brain size. Second, GNRs increase with brain size at the same rate in different circuits. Importantly, glia densities differ between regions, with the anterior piriform cortex having the highest density, followed by the posterior PCx, the neocortex, the cerebellum, and, lastly, the entorhinal cortex. The architectural principles identified in this study could be used as a means of discriminating brain circuits and gaining a better understanding of the extent to which neuronal function is influenced by glia.

## Materials and Methods

We used methods that were similar to earlier studies (Carlo and Stevens, 2013; Srinivasan and Stevens, 2018a, 2019). A previous study employing this approach showed that in contrast to neurons, the overall density of neocortical glia (with Nissl stains) is constant in the neocortex across species (Carlo and Stevens, 2013). In this study, we extend this approach to other brain regions such as the piriform cortex (shown in Fig. 1a). To validate the use of Nissl stains, we took advantage of available human and mouse tissue to examine counts of individual glial-types using anti-DAB (DAB: 3,3′-Diaminobenzidine) staining (Figs. 2, 5) as previously done (García-Cabezas et al., 2016), and compared them with counts obtained on tissue that were Nissl stained. We could not extend DAB staining to other species due to lack of available tissue, and used Nissl stains, instead. In this regard, the use of Nissl stains with stereological methods, combining light microscopy and morphological criteria, are still among the most efficient, especially when it is necessary to differentiate and count large quantities of glia and neurons across the layers of different brain regions and the brains of multiple species. Below, we present the stereological and histological procedures used.

**Figure 2:**
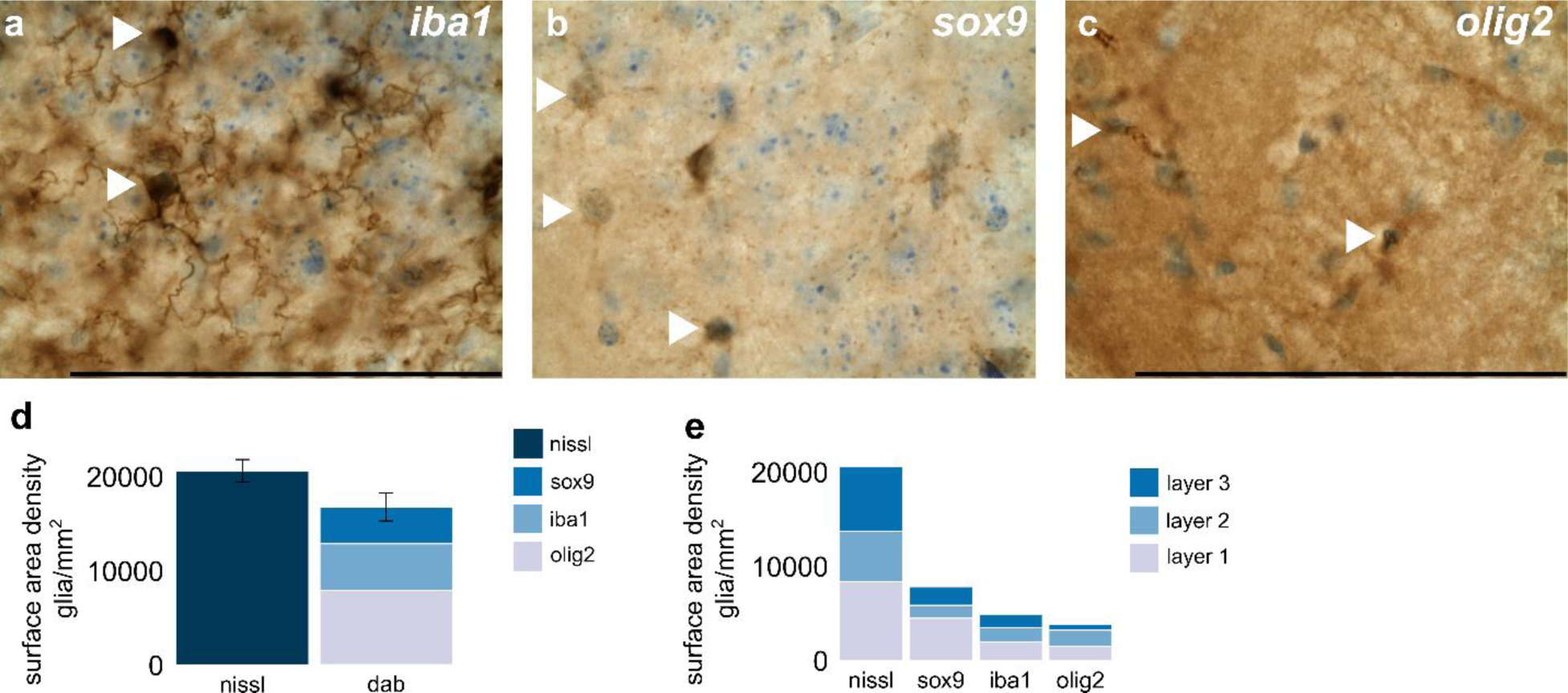
Glia in the anterior piriform cortex of mice. (a-c) high magnification images of the anterior piriform cortex in a mouse stained with DAB for marking astrocytes, microglia, and oligodendrocytes. In each image, the glia types can be distinguished (marked by arrows) by cells stained with DAB in brown, co-stained with Nissl in blue. (a) Microglia marked with Iba1 (b) Astrocytes marked with Sox9, and (c) oligodendrocytes marked with Olig2 In (a-c), white arrowheads highlight glial cell types co-stained with DAB and Nissl. (d-e) Estimates of number of cells under a mm^2^ of anterior piriform cortex using DAB and Nissl staining. (d) Shows the estimates of the total number of glia cell types measured with DAB and Nissl stains. The DAB staining bar is broken into three sections, each section a different color denoting the three major glia-types: sox9 in grey for astrocytes, iba1 in light blue for microglia, and oligodendrocytes in dark blue for oligodendrocytes. The bar on the right denotes the estimate of total number of glia under a mm^2^ of surface measured in Nissl stains. (e) Shows overall glia and each glial cell type distribution across layers. The Nissl bar shows the total number of glia in every layer, and the other three show the individual distribution of each glia-type in every layer. Scale bar: 100µm. See Figure S4 for the distribution of glial-cell types in each layer. Error bars in (d) are SEM.

### Histological procedures

We analyzed brain tissue from healthy adult specimens belonging to the following species: *Mus musculus* (mouse), *Rattus novergicus* (rat), Cavia porcellus (guinea pig), *Mustela putoris furo* (ferret), *Monodelphis domestica* (short-tailed opossum), *Felis catus* (domestic cat), and *Homo Sapiens* (humans). Animal care protocols were approved by the Salk Institute Animal and Use Committee and conform to US Department of Agriculture regulations and National Institutes of Health guidelines for humane care and use of laboratory animals. Each specimen was perfused with aldehyde fixative agents and stored long-term in 10% formalin. In preparation for cutting, all brains were submerged in solutions of 10% (wt/vol) glycerol and 10% formalin until the brain sank, and then moved into 20% glycerol and 10% formalin until the brain sank again; the average time in each solution was 3 to 10 d. These cryoprotected brains were then cut coronally on a freezing microtome at a thickness of 40 or 50 μm. Every 6^th^ section was stained with thionin for visualization of Nissl bodies.

### DAB/Nissl staining

To ascertain that glia estimates from Nissl stains across species are accurately representative of glia numbers, we stained for the three major types of glia (separately) -- astrocytes, microglia, and oligodendrocytes – and compared the cumulative number of glia estimated through this method with glia estimated from Nissl stains. A previous study has established the effectiveness of this approach (García-Cabezas et al., 2016). We undertook the comparison estimations for mouse and human sections from the anterior piriform cortex region, but not for other species, as that tissue was not available to us.

The procedure for obtaining tissue slices (50 um sections) is described in Histological procedures. For the first step of DAB staining, we washed the sections in 10 ml 0.1M PO_4_ buffer with 1 ml 30% Hydrogen Peroxide (H_2_O_2_) for 10 minutes, three times, to reduce non-specific peroxidase reaction. We then incubated the sections in a blocking solution comprising 0.1 M PO_4_ buffer, 3 % normal animal serum (up to 5%), and 1 % of 25% Triton-X for 1-4 hours at room temperature on a rotator, to block non-specific antigen reactions and to prepare solutions for primary antibody incubation. We then incubated sections in blocking solution with primary antibodies at the prescribed concentrations (given below) overnight at 4 degrees on a rotator. The next day, we incubated sections in blocking solution along with the biotinylated secondary antibody at 1:200 for 1-2 hour(s) at room temperature on the rotator. We washed the sections three times, for 10 minutes each time, on a rotator at room temperature. We then incubated the sections in ABC (Acetyl-Avidin Biotin Peroxidase Complex) solution for 1-2 hours at room temperature. The ABC solution was prepared using the Elite ABC Kit (Vector Laboratories # PK-6100), wherein 1 drop of A solution was mixed with 1 drop of B for every 5 ml of 0.1 M PO_4_ buffer, then left standing for 30 minutes before use. The sections were then washed three times for 10 minutes each at room temperature on the rotator. After the washes, the sections were treated to the DAB solution. The DAB solution was prepared by dissolving a DAB tablet (10 mg) in 30 ml 0.1 M PO_4_ buffer and filtered with rapid flow filter paper. We then added 33 µl of 30% H_2_O_2_ to the filtered DAB solution and mixed the solution well. We transferred sections to this solution in a wide petri dish and manually swirled each section until it turned rusty brown. Finally, the sections were washed again in buffer three times. Following this procedure, we mounted the sections on glass slides and stained the sections for Nissl as described in the Nissl Staining section.

We used the primary and secondary antibodies at the following concentrations. To mark astrocytes, we used the Sox9 anti-mouse antibody (1:1000, abcam, ab185230) and secondary biotin conjugated anti-mouse at 1:250 (ebioscience, cat: 13-4013-85). To mark microglia, we used the Iba1 anti-donkey antibody at 1:500 (FUJIFILM Wako Pure Chemical Corporation, 019-19741) and secondary Biotin-SP-conjugated AffiniPure Donkey Anti-Rabbit IgG (H+L) at 1:250 (Jackson ImmunoResearch cat: 711-065-152). For oligodendrocytes, we used the Olig2 anti-mouse antibody (1:100, abcam ab109186) and secondary Biotin-SP-conjugated AffiniPure Donkey Anti-Rabbit IgG (H+L) at 1:250 (Jackson ImmunoResearch cat: 711-065-152).

### Nissl staining

The tissue was defatted with 100% Ethanol: Chloroform (1:1) overnight, rehydrated in a decreasing alcohol (with DI H_2_O) series (100 %, 95, 70, 50), then treated with DI H 2 O, Thionin stain, followed by DI H_2_O, an increasing alcohol series (50 %, 70, 95, 95, 100, 100), Xylenes I, Xylenes II, and then cover-slipped. The tissue was dipped 4-5 times in each of the solutions for 1 minute except for the thionin stain (1-2 minutes), and Xylenes II (1 hour). The thionin stain was prepared by first gently heating 1428 ml DI H2O, 54 ml of 1M NaOH, 18 ml of glacial acetic acid until the solution was steaming. Then, 3.75 g of thionin was added and the solution was boiled gently for 45 min, cooled, filtered, and used for staining.

### Volumetric estimate

The layers of the anterior piriform cortex, the posterior piriform cortex, and entorhinal cortex, were outlined in successive coronal sections across the rostral-caudal axis with the help of Neurolucida (version 10.53; MBF Bioscience, Wilmington, VA) at low magnification (2X and 4X objectives) based on standard atlases and primary literature (Snider and Niemer, 1961; Neville and Haberly, 2004; Paxinos and Franklin, 2004; Mai et al., 2015; Ding et al., 2016). Once the layers from each individual section were outlined, all sections were aligned for subsequent three-dimensional reconstructions with Neurolucida explorer. Alignment was done by starting with the second section and aligning each section to the previous one. Surface areas and volumes for the anterior piriform cortex were then obtained from three-dimensional reconstructions.

### Glial density

10- or 20-μm-wide columns, perpendicular to the pial surface, extending down to the boundary of the deepest layer, were delineated with the Neurolucida contours option. The sections and columns within them were chosen so that we had equal coverage of the region along the rostral-caudal and dorsal-ventral axes. For obtaining surface area density counts (number of glia/mm^2^), three measures were used: the width of the column, the thickness of the section, and the number of neurons in the column. Columns were randomly chosen while ensuring that they were close to perpendicular to our coronal cut and the surface of the brain (as an illustration, Table S4 shows the number of columns in each species for APCx). Glial cells were manually counted with standard unbiased stereology techniques in Nissl-stained sections at 100X oil magnification (Fig. 1b), using Neurolucida 10.3 (MBF, Vermont, USA). The counting column functioned as a typical dissector whereby neurons and glia were marked as they came into focus given that they also fell within the acceptance lines of the dissector.

Glia were differentiated from neurons based on size (smaller) and morphology (more punctate versus distinctive shape and processes extending out for neurons) of the cell, and by the presence of a distinct nucleolus in neurons (Fig. 1b). Previous studies have shown that even though astrocytes have large cell bodies, in Nissl stained sections the cellular size that is stained is significantly smaller than neurons (Jyothi et al., 2015; García-Cabezas et al., 2016). Cells that were hard to distinguish (less than 10 %) were labeled as unknown and not included in the counts. Approximately 50 to 100 objects of interest were counted in each 3D column. We counted all the glia in our column from the pial surface to the white matter without the use of guard zones. A detailed discussion of guard zone use, as it pertains to stereological methods and data collection for frozen sections, is reported in (Carlo and Stevens). Although it was outside of the scope of this study to explain differences between cell layers, it is noteworthy that the number of glia (Figs. 1a, S1 and Table S1) and astroglia, in particular (see section 2.3 below, Figs. 1d, S1), was different across layers.

### Astrocytes in SOX9-EGFP mice

To test if astroglia, in particular, showed a similar trend as glia as a whole, we examined APCx in brains of SOX9-EGFP male mice (∼4 months old, fixed in 4% PFA and postfixed in the same solution for 24 h), which were generously donated by Dr. Scott Magness (University of North Carolina at Chapel Hill). The tissue was kept at 4°C in 30% sucrose/PBS solution until flash-frozen using optimal cutting temperature medium (ThermoFisher), and cut into 40-70 μm sections on a cryostat. Sections were washed with PBS before being incubated for 4h in a PBS-based blocking solution containing 10% normal goat serum (NGS), 0.1% Triton X-100 (Sigma-Aldrich) and sodium azide. Sections were then incubated overnight at 4°C in PBS containing 2% normal goat serum (NGS), rabbit anti-GFP Alexa Fluor 488-conjugated (1:500) and mouse anti-NeuN (1:250) primary antibodies. Sections were then washed with PBS and incubated with anti-mouse Alexa Fluor 568-conjugated secondary antibody (1:1000) and then mounted in DPX mounting medium. Images were collected with an LSM 880 inverted Zeiss confocal microscope.

Randomly chosen columns, perpendicular to the pial surface, and containing layers 1 to 3 were delineated using Neurolucida 10.53. For the quantification of astrocytes, the SOX9 channel (568 nm) was made invisible and the number of GFP+ cells observed at 20x magnification was counted for each layer using Neurolucida 10.3, MBF, Burlington, Vermont. The GFP (488 nm) channel was then made invisible and SOX9+ cells were counted using the same methodology. The counting column was drawn in a single section 50-100 µm thick located 15-30 µm deep from the top of the slice. Note that NeuN staining intensity did not match the endogenous GFP staining across the thickness of the tissue. For this reason, we chose to count astrocyte GFP marked cells and neurons in the slice, where they were both stained equally well.

### Statistical analysis

For the plots in Figures 3--6 and S2--5, we obtained the Pearson’s correlation coefficient using the R statistical programming language. We used inbuilt functions from R for calculating the best line fit, correlation coefficient, and the 95% confidence interval by fitting the data to a linear model. We have listed the correlation coefficient in the figures and the results section of the main text.

**Figure 3:**
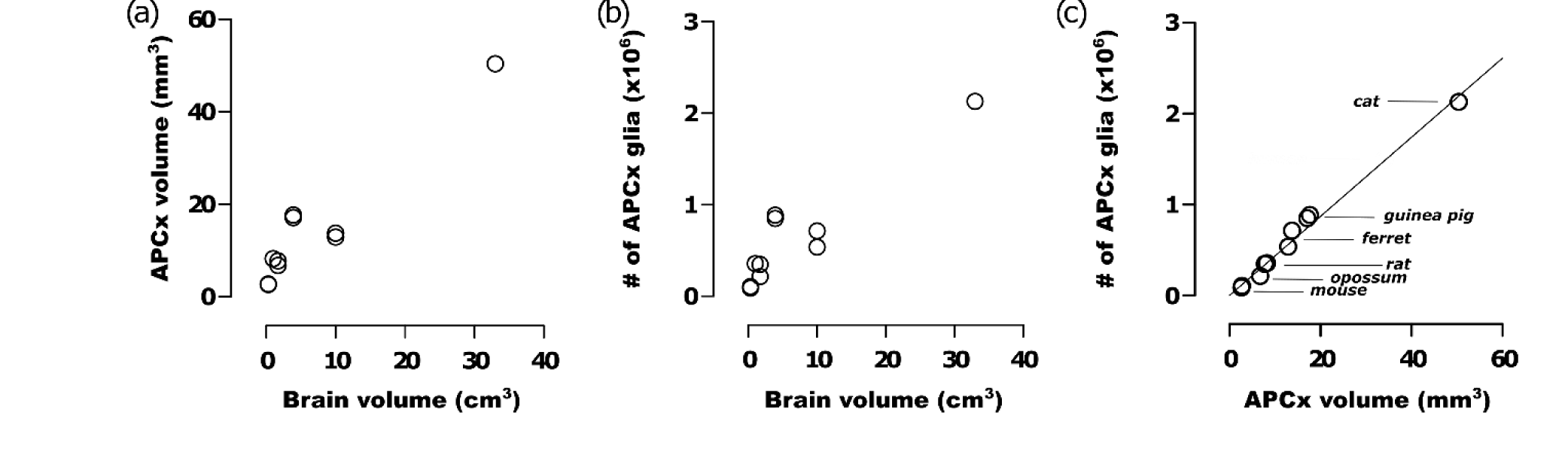
Glia volume densities were constant across species in the APCx. The APCx volume (a) and number of glia contained in it (b) increase with brain volume. (c) Despite significant differences in APCx volume and absolute number of glia across the different species studied, the number of glia per mm^3^ was approximately the same, being an excellent fit for the equation, y = 43,500x with an R^2^ of 0.98. The species examined were mouse, rat, opossum, guinea pig, ferret, and cat, in the order of increasing brain volume. See also Tables 1 and 2 for the numerical values plotted in the graphs, and Figures S3d for a comparison of densities between regions.

**Table 1.** Summary of APCx measurements. Values correspond to the data points in Figure 2. For animals with more than one specimen, we list averaged values. For individual specimens’ measurements, see Tables S1 and S2.

| Species' common name | Brain volume (cm<sup>3</sup>) | APCx surface area (mm<sup>2</sup>) | APCx Volume (mm<sup>3</sup>) | APC width (mm) | Surface density (x 1000 glia/mm<sup>2</sup>) | Number of glia (x 1000) |
| --- | --- | --- | --- | --- | --- | --- |
| Cat | 33 | 58.0 | 49.9 | 0.88 | 37.5 | 2175 |
| Human | 1300 | 32.4 | 34.02 | 1.05 | 46083 | 1493 |
| Ferret | 10 | 19.4 | 13.2 | 0.69 | 31.5 | 611 |
| Guinea pig | 3.9 | 19.8 | 17.2 | 0.85 | 43.5 | 862 |
| Mouse | 0.3 | 4.7 | 2.6 | 0.55 | 18.0 | 85 |
| Opossum | 1 | 9.5 | 8.3 | 0.86 | 37.4 | 355 |
| Rat | 1.7 | 10.7 | 7.2 | 0.68 | 28.0 | 298 |

**Table 2.** Summary of PPCx and ECx measurements. Values correspond to the data points in Figure 3.

| **Species' common name** | **Brain volume (cm<sup>3</sup>)** | **PPCx width (mm)** | **ECx Volume (mm<sup>3</sup>)** | **Number of glia under 1 mm<sup>2</sup> (x 1000)** | **Number of glia in ECx (x 1000)** |
| --- | --- | --- | --- | --- | --- |
| **Cat** | 33 | 0.884 | 60.4 | 21.2 | 2175 |
| **Ferret** | 10 | 0.86 | 19.2 | 26.4 | 611 |
| **Mouse** | 0.3 | 0.45 | 2.36 | 16.9 | 85 |
| **Rat 1** | 1.7 | 0.685 | 8.59 | 16 | 298 |
| **Rat 2** | 1.7 | 0.55 |  | 14.6 |  |

For the density counts, assuming a homogenous spatial distribution, the column density counts should approximate a Gaussian distribution. This happens because with a spatially homogenous distribution, as one traverses a line from a fixed point, one will encounter glial cells. The probability of encountering a glial cell is given by a Poisson distribution in this case, as their distribution is homogenous. In the limit, with a large number of cells, the distribution of densities of different columns will follow a Gaussian distribution. We validated our column counts by fitting a relative cumulative distribution of glial cell densities across columns to a cumulative Gaussian and testing the count distribution with a Shapiro-Wilks test (W > 0.05).

## Results

We had two major goals in this study: first, to examine if the volume density of glia (i.e., number of cells per mm^3^) or number of glia relative to neurons changed within a brain region across mammalian species, and second, to express those relationships quantitatively. To achieve a broad representation of the brain, we collected data from the anterior piriform cortex, the posterior piriform cortex (both of which are part of paleocortex - a subtype of allocortex), the entorhinal cortex (part of the periallocortex – also a subtype of allocortex), and reanalyzed data available in the literature for the neocortex and the cerebellum. We started by investigating if, as for the 6 - layered neocortex, glia occupied a constant volume fraction of the thinner 3-layered paleocortex.

### Estimates of Glia in anterior piriform cortex with Nissl and DAB staining are similar

Our first task, since we only had Nissl-stained tissue for all species analyzed, was to ascertain that glia count measures estimated using Nissl stains are accurate. To do so, we used two methods to estimate the number of glia in human (shown in a later section) and mouse anterior piriform cortex. With the first method, we used Nissl stains, and in the second, we marked three types of glia – astrocytes, microglia, and oligodendrocytes – simultaneously with Nissl and anti-DAB staining, as done previously (García-Cabezas et al., 2016). For instance, we marked astrocytes by staining for sox9 using DAB, and then co-stained for Nissl (using Thionin). We then manually counted the number of astrocytes by only marking cells that had simultaneous expression of DAB and Nissl, as shown in Figure 2b. We repeated this procedure for microglia (marked with Iba1, Fig. 2a), and oligodendrocytes (marked with Olig2, Fig. 2c). We independently validated our DAB staining measures for astrocytes by also estimating the number of astrocytes in a sox9-GFP mouse (Fig. 1b, c). We found that the estimate of astrocytes to neurons using the sox9-GFP mice and DAB staining for sox9 in normal mice was nearly identical (Fig. S1). Next, we present our estimates of glia with Nissl and DAB staining.

We found that the estimates of glia obtained by Nissl and DAB staining are comparable (Fig. 2d). We highlight the mouse findings here, and human findings in a later section (Fig. 6). The counts in Figure 2d are presented, for all glia (Nissl dark blue bar – Nissl stains) and individually for each glia type (dab bar: grey – astrocytes, light blue – microglia, medium blue – oligodendrocytes). The estimates of how total glia (from Nissl stains) and individual glia-types (from DAB staining) are allocated to each layer is presented in Fig. 2e. Figure 2d shows that the counts obtained through both measures are similar, with DAB staining yielding a lower estimate. The lower estimate, most likely, arises from two sources. First, we only estimated the three major glia cell types, and not minor types like ependymal or radial glial cells. Second, studies have reported that the marker for oligodendrocytes, Olig2 (Valério-Gomes et al., 2018), that we used, does not always capture all oligodendrocytes(Huang et al., 2022). The lower estimate, thus, is a reasonable measure of total number of glia. The match gave us confidence that we could use Nissl stains to estimate glia densities across species, and we present these estimates in the next section. But, first, we examine the composition of glia across layers starting with astrocytes.

Astrocytes are important for synaptic physiology, transmission and plasticity (Verkhratsky et al., 2015; Allen and Eroglu, 2017; Allen and Lyons, 2018). The higher numbers of synapses in Layers 1 and 3 compared to Layer 2 (Neville and Haberly, 2004) suggests that we should observe more astrocytes in Layers 1 and 3. Analysis of glia as a whole supports this view as Layer 1 contained most of the glia, followed by Layer 3. Layer 2, which is particularly neuron-rich (Figs. 1a, b, S5a) contained the fewest glia. When we inspected astrocytes by themselves, indeed, this was case, too (Figure 1e). We used an alternative approach to validate our astrocyte results with DAB staining. We examined the number of astrocytes using a Sox9-GFP mouse, in which astrocytes are specifically labeled with GFP (Sun et al., 2017). As described in the methods section, Figure 1 shows coronal sections through the piriform cortex, with Sox9-positive cells marked in green, and NeuN-positive cells (denoting neurons) marked in red. Figure S1 shows two comparisons. First, it shows the comparison between astroglia and neurons across the three layers with the highest number of astroglia (Fig. S1a) in layer 1. Second, it highlights the similarities in astrocyte/neuron ratio obtained through either the Sox9-GFP mouse or DAB staining suggesting that DAB staining is effective in estimating astrocyte number.

With regards to the other glial cell types (Fig. 1e), microglia showed a trend similar to astrocytes with most of them located in Layer 1, followed by Layers 3, and 2. Oligodendrocytes, however, deviated from this trend, being equally high in Layers 1 and 2. Notably, astrocytes were the biggest proportion of glial cell types in Layers 1 and 3, which contain most of the glia, and thus, were the most prevalent glial cell type. Overall, these features of glia distribution (Figs. 2e, S5b) might showcase the computational processes carried out in each layer (Suzuki and Bekkers, 2011; Bekkers and Suzuki, 2013a).

### The volume density of glia is conserved in both parts of the piriform cortex across brain sizes

A previous study (Carlo and Stevens, 2013) showed that glia densities are conserved in the motor, somatosensory, parietal, and temporal regions of the neocortex across four mammalian species (21,100 glia/mm^3^). An even earlier study (Stolzenburg et al., 1989) found a similar conservation of glia in the neocortex in five insectivore species (Fig. S2, 25,600 glia/mm^3^, CI: 24,000 – 27,200). To test if glia densities are conserved across regions and species, we examined other regions including the anterior piriform cortex (APCx) and posterior piriform cortex (PPCx).

We estimated the total number of glia in the anterior piriform cortex (APCx) of six mammalian species (Fig. 3) and determined how they varied with APCx size, measured in terms of volume and surface area (Fig. 3a-c, Fig. S2). We found that the number of glia increased proportionally with APCx volume (Fig. 3c). The relationship was an excellent fit for the equation *N_G_* = *m*\**V_APCx_* (R^2^ = 0.98, CI: 39.89 – 48.09, *N_G_* = number of glia, *V_APCx_* = volume of APCx), where the slope *m* = *N_G_/V_APCx_* represents the volume density of glial cells (*G_v_*), which was found to be 43,500 glia/mm^3^ (95% confidence interval: 39,890 – 48,090 glia/mm^3^). The number of glial cells was also proportional to the surface area of APCx (Fig. S2).

To validate our calculations, we used additional ways of measuring volume densities. First, we estimated glial volume density by measuring the surface density of glia (number of glia under a mm^2^ of APCx, Figure S2b and also Table 1, Table S1 for individual animal counts), and the thickness of the APCx (Fig. S2c, Table 1, and Table S1 for individual animal counts), similar to earlier studies (Carlo and Stevens, 2013). Surface area density and thickness are related by the equation *G_sad_* = *N_0_* + *m*\**T_APCx_*, where *G_sad_* is the surface density in glia/mm^2^, *N_0_* is an additive constant (intercept) in units of glia/mm^2^, *T_APCx_* is the thickness of the APCx, and *m* is the slope in units of glia/mm^3^ (glial volume density). The data were fit to this equation (Fig. S2d). We found that the regression line had an R^2^ value (coefficient of determination) of 0.89, and the volume density of glial cells (*G_v_*) was 55,000 glia/mm^3^ (95% Confidence Interval, 24,000-99,000). With the next measurement methodology, we directly estimated the average glial volume density from our measurements of cortical columns across the six species and found glia density to be 44,249 ± 2,140 glia/mm^3^ (mean ± sem, Table S3), which is not significantly different from the estimates obtained from Figure 3c. Thus, for each glial cell that is added to the piriform cortex, neuropil volume was increased by an amount that covered 0.022 nL (1/volume density).

Ass described above, the calculations of glia density using different methodologies were not significantly different from each other. Therefore, for brain regions where we did not have estimates for the volume of the region, such as PPC that we describe in the next section, we used these methods.

Unlike APCx, PPCx is marked by the absence of the Lateral olfactory tract or LOT (contains olfactory bulb nerve input to PCx) and a lower neuronal density (Srinivasan and Stevens, 2018a), in addition to having significantly less input from the olfactory bulb and more associative input (Neville and Haberly, 2004). Further, PPCx contains surface-associated astrocytes (SAAs), which are unique to this brain region, given their direct apposition to the cortical surface and large-caliber processes that descend into layer 1 (Feig and Haberly, 2011). Functional studies in humans and rodents have suggested that PPCx function differs from that of APCx, mirroring their morphological differences (Haberly, 2001a; Calu et al., 2007; Srinivasan and Stevens, 2018b). We, therefore, asked whether, despite the proximity between the two regions, glia densities were different between them.

We observed that glia volume density was lower in PPCx compared to APCx (Fig. S3c vs. Fig. 3c). We found that, once again, glia occupied a constant volume density in PPCx of the four species analyzed (Fig. S3). As Figure S3 shows, as the width of the PPCx increases with increasing brain size, so does the number of glia under a square mm. We obtained volume density by normalizing surface area density to PPCx width and found it to be 27,750 glia/mm^3^ (CI: 24,320 – 31,180). The lower density of glia in PPCx compared to APCx reflects both the lower density of neurons in PPCx, the lower number of olfactory bulb inputs as well as synapses in layer 1. Thus, the concomitant decrease in glia density with neuronal density and number of synapses suggests glia quantity are an additional criterion underlying morphological and functional differences between the two regions.

### Glia volume density in the entorhinal cortex remains constant with increasing brain sizes

Next, we investigated if the conservation of glia volume density extended to an architecturally and functionally distinct region: the periallocortex. We examined the entorhinal cortex, a periallocortical region that processes inputs from topographic (e.g., visual) and distributed (e.g., olfactory) circuits. The entorhinal cortex (ECx) is a multimodal circuit that closely interacts with the hippocampus (Witter et al., 2017) and is integral to spatial navigation (Fyhn et al., 2004). In this sense, it differs from many of the neocortical regions examined in earlier studies and from the anterior and posterior segments of the piriform cortex, as these regions are primarily sensory regions in which sensory input from peripheral organs (e.g., nose and eyes) is processed.

As Figures 4a—c show, ECx volumetric and glia measurements show a trend comparable to that of olfactory regions. Glia occupy a constant fraction of volume in the entorhinal cortex (Fig. 4c). Their number is proportional to the volume of the entorhinal cortex as described by the relation, N_G_ = 11,000*Vol_ECX_ (R^2^ = 0.969, CI: 7,490 –12,570; glia density = 11,000 glia/mm^3^).

**Figure 4:**
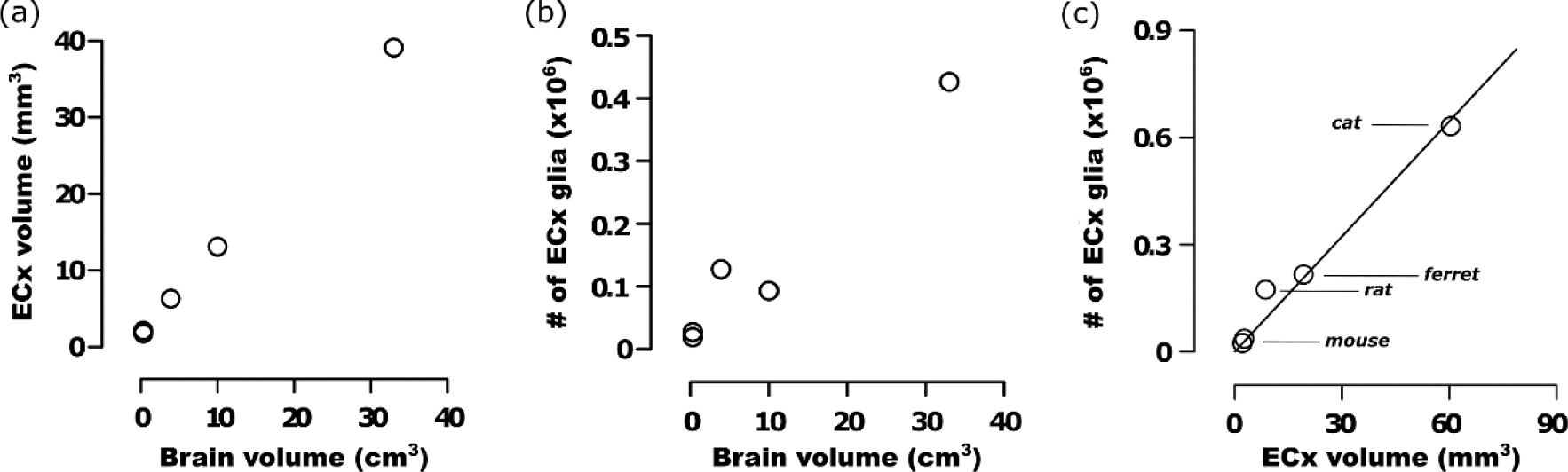
Glia volume densities were constant across species in ECx. The ECx volume (a) and number of glia contained in it (b) increased with brain volume across the species studied: mouse, rat, ferret, and cat. (c) The number of glia per mm^3^ was approximately the same in the ECx across the species studied (mouse, rat, ferret, and cat), and was fitted with a line represented by the equation y = 11,000x with an R^2^ of 0.95. See also Tables 1 and 2 for the numerical values plotted in the graphs, and Figures S3d for a comparison of densities between regions. The species examined were mouse, rat, opossum, guinea pig, ferret, and cat, in the order of increasing brain volume.

Taken together, these data suggest that, for any given region, the number of glia per unit volume remains constant even as brain size changes. Notably, our data also shows that the volume density of glia is distinct for APCx, PPCx, and ECx (Fig. S4d). Note that the confidence intervals for glia density do not overlap.

### The number of glia per neuron increases with bigger brains in the paleocortex and prefrontal cortex

A previous study of the somatosensory, motor, parietal and temporal regions of the neocortex showed that while neuronal density decrease with bigger brains, glia densities remain constant, implying that glia to neuron ratios increase (Carlo and Stevens, 2013). To test if this trend might be general, we estimated glia-neuron ratios (GNR) and their relation to brain size in brain regions besides the neocortex. In the piriform cortex, glia-neuron ratios increase with brain size, best described by a power-law equation: glia/neurons = 0.7\**b_v_*^1/4^ (Fig. 5a), where *b_v_* is brain volume. The data for neurons in Figure 5a is from a previous study (Srinivasan and Stevens, 2019), while the data for glia is from this study. Note that in a power-law relationship, the dependent variable *y* is related to the independent variable *x* according to the equation *y* = a*x*^b^, where a and b are constants. b is called the scaling exponent and, when positive, determines how fast *y* grows with respect to *x* (with negative b’s, the equation captures how fast *y* reduces with *x*; b=1 implies that *y* grows proportionally with *x*).

**Figure 5:**
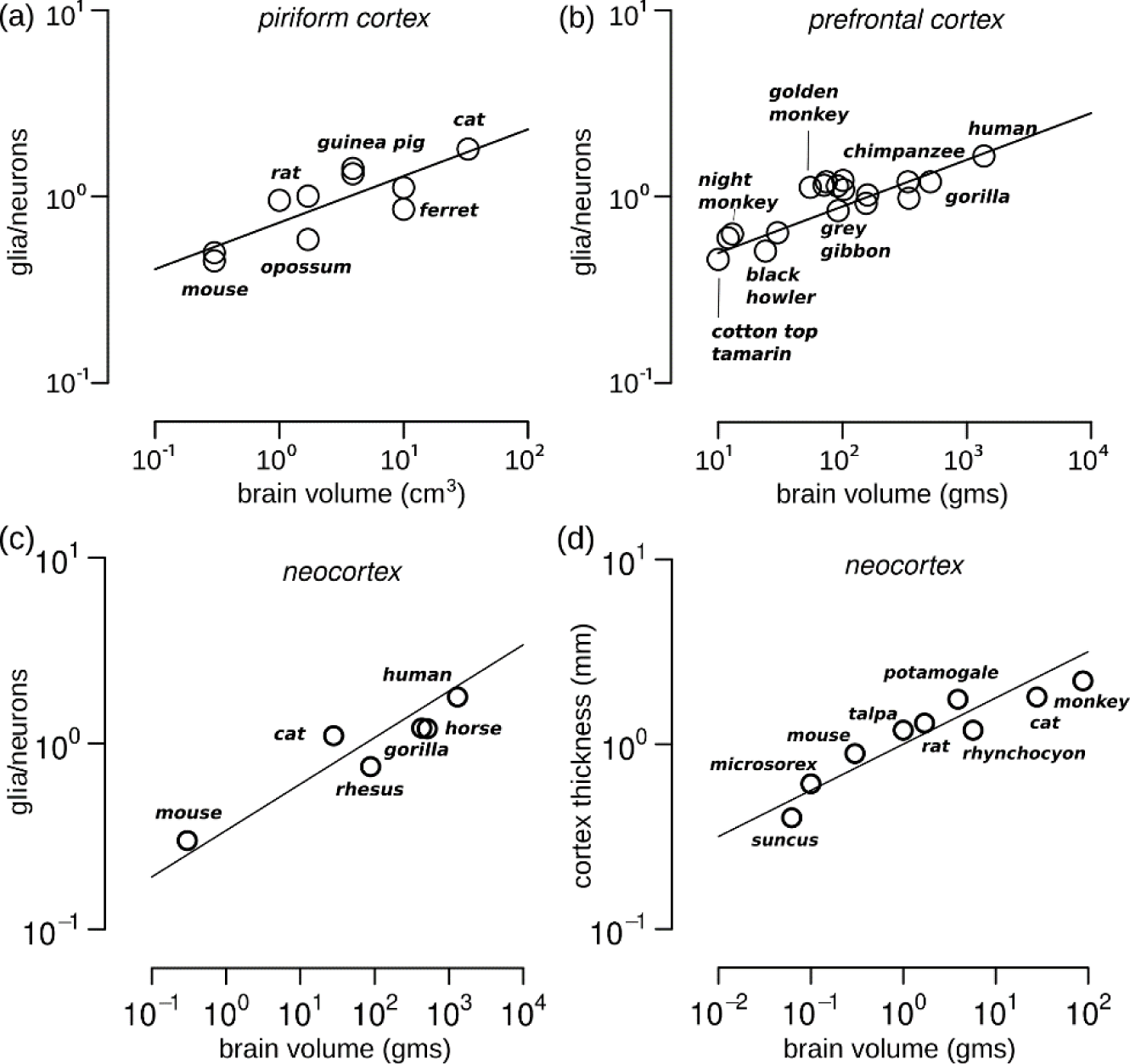
Glia-neuron ratio increased with the size of the brain. (a) GNR in APC of six mammals (mice, rats, opossums, guinea pigs, ferrets, and cats) shows that in APC too, GNR increased with brain size as y = 0.72x^1/4^ with an R^2^ of 0.67 and CI of 0.1-0.38. Note that most species have multiple animals that were examined. For number of glia, see Table 1, and for number of neurons see Table S3 of (Srinivasan and Stevens, 2019) (b) GNR replotted from (Sherwood et al., 2006) showing that as the size of the primate brain increases, GNRs in PFC increase. The numerical values were fitted with a line according to the equation y = 0.3x^1/4^ with an R^2^ of 0.68 and a CI of 0.14-0.29. (c) GNR in the neocortex as a whole measured in 6 mammals shows that as brain volume increases across species, so does the ratio of glia to neurons. The data are fit to a power-law equation y = 0.34x^1/4^ with an R^2^ of 0.88.(d) The thickness of the cortex increases with brain size, represented here with weight in grams. The data includes 5 insectivores (Stolzenburg et al., 1989) and 4 mammals (Carlo and Stevens, 2013). The fitted line has the equation, y = 0.34x^1/4^, with an R^2^ of 0.86.

We also examined regions that were architecturally and functionally different, to test if there were similar trends. In a previous study, Sherwood and colleagues (Sherwood et al., 2006) reported a similar increase in glia-neuron ratios with brain size in the prefrontal cortex of primates. Remarkably, when we replotted the data from the study (Fig. 5b), we found that, here, too, glia-neuron ratios increased with brain size according to a power-law: glia/neurons = 0.3* *b* ^1/4^, where b_V_ is the brain volume. Interestingly, a 1/4 power-law relationship between glia-neuron ratios and brain volume was also observed in the cerebellum for the ratio of Purkinje cells to glia in the molecular layer, where parallel fibers synapse onto Purkinje cells: glia/neurons = 8.75* *b_V_* ^1/4^ (Fig. S4c, data from (Friede, 1963)). The brain volumes in this plot, as well as in Figure 5a, were taken from (Haug, 1987), and, for the sake of consistency, we only considered glia-neuron ratios for the cerebellum of those animals whose brain volumes were included in the Haug study. In summary, GNR scale with brain size similarly in APC, prefrontal cortex, and the cerebellum.

Finally, the conserved GNRs in these three regions: PCx, PFC, and Cerebellum led us to wonder if this might extend to the cortical region as a whole. We examined the literature (Haug, 1987; Christensen et al., 2007; Verkhratsky and Butt, 2018) and found glia to neuron ratios in 6 mammals, including humans, for the whole neocortex and shown in Figure 5c. Here, too, as brain volume increased, so did the number of glia per neuron according to a power-law glia/neurons = 0.34* *b* ^1/4^. Notably, the constant factor or the intercept in the log-log plot for the neocortex was not significantly different from that of PFC (Fig. 5b) and piriform cortex (Fig. 5c), suggesting that the scaling of the ratio of glia per neuron might be a common law across brain regions and species.

### Explanation for GNR power-laws

Why would GNR follow a ¼ power law? To answer this question, we examined data from the neocortex collected in two earlier studies (Stolzenburg et al., 1989; Carlo and Stevens, 2013). Carlo and Stevens showed two principles of conservation in mammalian neocortices: glial volume densities are the same (*g_vd_*: 21,000 glia/mm^3^) across species, and neuronal surface area densities (*n*_*sa*_: number of neurons under a mm^2^ of cortical surface) are the same (*n_sa_*: 100,000/mm^2^). From these two findings, we can express the number of glia and neurons under 1 mm^2^ of cortical surface as *glia* = *volume* * *g*_*vd*_ = *g*_*vd*_ * 1 * *t* = 21,000*t*, and *neurons* = *n*_*sa*_ * *surfacearea* = *n*_*sa*_ * 1 = 100,000, where *t* is the thickness of the neocortex. Furthermore, the thickness, *t*, (examined in (Stolzenburg et al., 1989; Carlo and Stevens, 2013) and Fig. 5d, covering rodents, carnivores, primates, and insectivores) can be expressed as a function of brain weight (*b_wt_*) by the equation, *t* = *b_wt_*^1⁄4^. Thus, GNR is given by

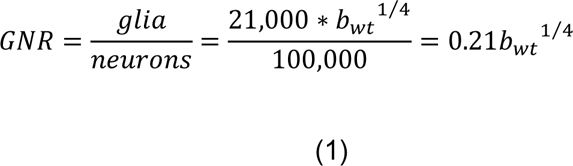

The result above shows that the GNR power-law arises because glial volume densities are conserved, and so the number of glia depend on the thickness of the cortex, which varies with brain volume as a ¼ power-law (Fig. 5d). Thus, collectively, these results, which show glia-neuron ratios scaling with brain volume according to a ¼ power law, independent of brain region, architecture, and function, suggest a common principle of brain organization.

### Humans have a higher number of glia but a similar volume density to mice

Previous reports have pointed out that humans might have a higher number of glial cells than other species (Diamond et al., 1985). To test if this might be true, we estimated the number of glia in the anterior piriform cortex of humans. Similar to mice, we estimated total number of glia using Nissl stains, and numbers of individual glial types – astrocytes, microglia, and oligodendrocytes – using DAB staining.

When using Nissl stains in humans, we estimated the total number of glia under a square mm of human anterior piriform cortex (APCx) to be 46,083 (Fig. 6). The surface area of the human APCx was 32.4 mm^2^ showing that there are around 1.5 million glia in the human APC. By comparison, the total number of glia in the mouse APC was 80,000; an order of magnitude lower than humans. The greater number of glia in humans results from human PCx being much larger by volume partly because the surface area of PCx is larger, and also because PCx is thicker reflected in human PCx having a higher surface density of glia than mice (at 20,000) or species with larger brains (than mice) such as cats (37,000).

**Figure 6:**
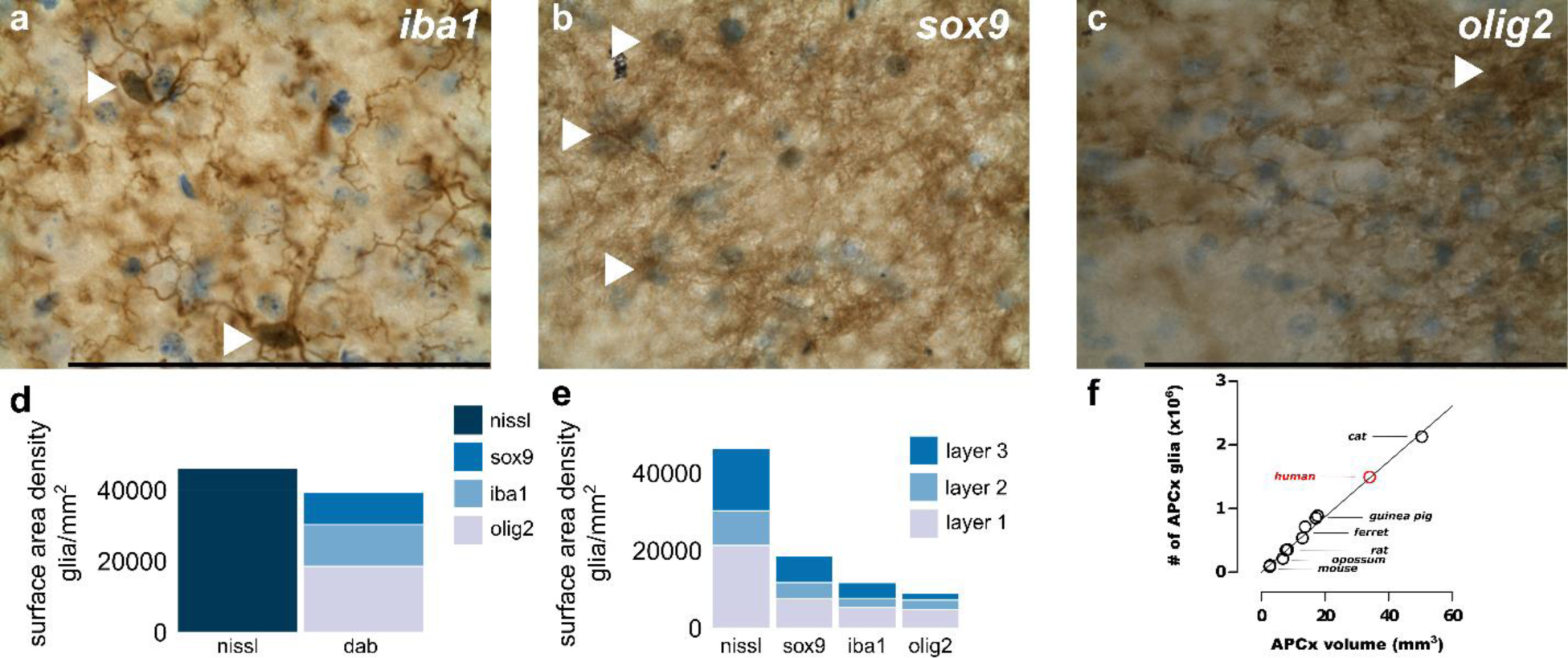
Glia in the anterior piriform cortex of humans. (a-c) high magnification images of the anterior piriform cortex in a human stained with DAB for marking astrocytes, microglia, and oligodendrocytes. In each image, the glia types can be distinguished (marked by arrows) by cells stained with DAB in brown, co-stained with Nissl in blue. (a) Microglia marked with Iba1 (b) Astrocytes marked with Sox9, and (c) oligodendrocytes marked with Olig2. (d-e) Estimates of number of cells under a mm^2^ of anterior piriform cortex using DAB and Nissl staining. (a) Shows the estimates of the total number of glia cell types measured with DAB and Nissl stains. The DAB staining bar is broken into three sections, each section a different color denoting the three major glia-types: sox9 in grey, iba1 in light blue, and oligodendrocytes in dark blue. The bar on the right denotes the estimate of total number of glia under a mm^2^ of surface measured in Nissl stains. (e) Shows the layer wise distribution of glia assessed. The Nissl bar shows the total number of glia in every layer, and the other three show the individual distribution of each glia-type in every layer. (f) Replotting the scaling of APCx volume versus number of glia shown in Figure 3c, with human data. The human data point (in red) falls on the regression line with slope 43,000 showing that evolution has preserved glial population densities across species. Scale bar: 100µm.

**Figure 7:**
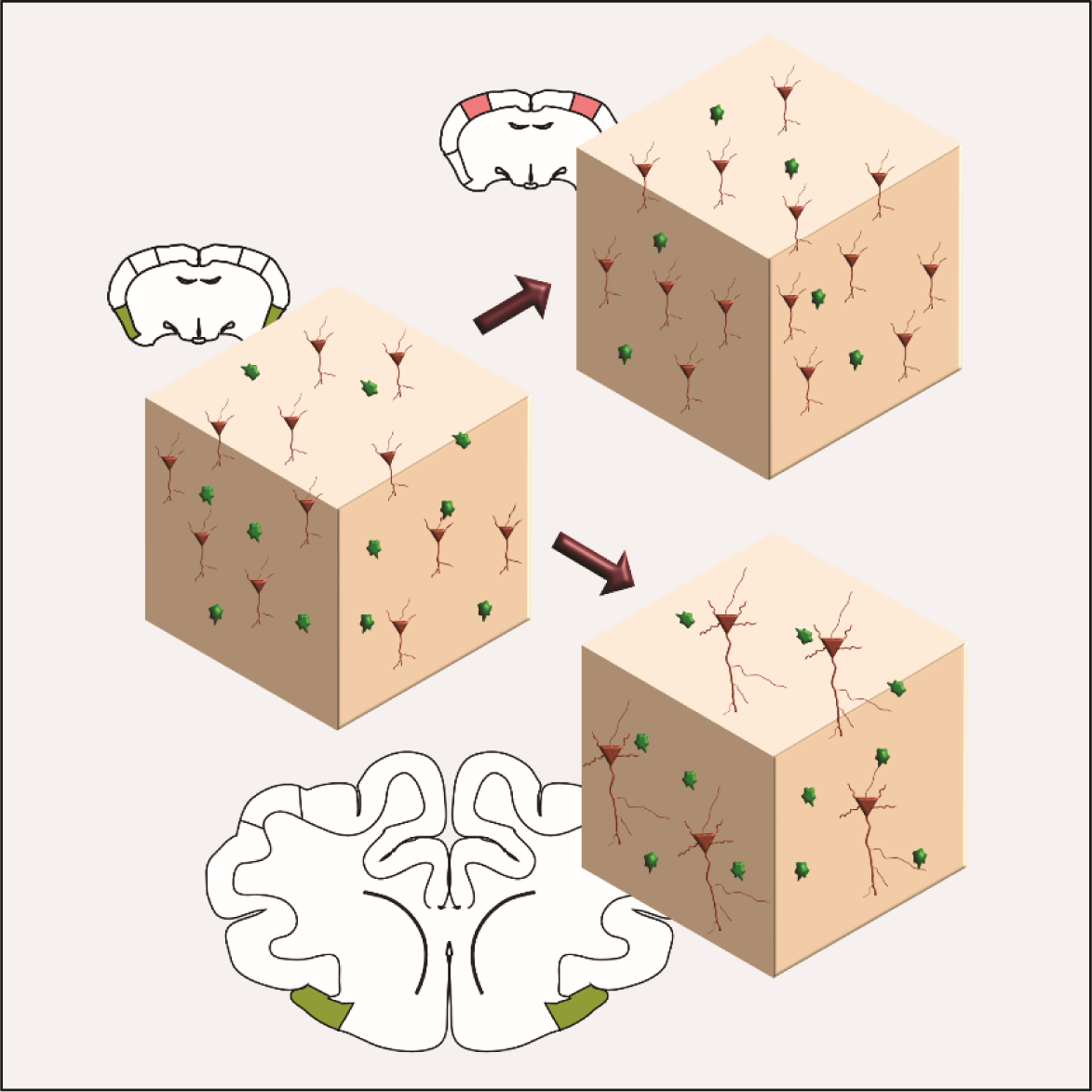
Two principles of glia organization. First, glia density is conserved within a brain region across species. Compare the piriform cortex, shaded in green, in two species (mouse and cat). Even as the piriform and brain sizes change between the two species, the number of glia is conserved within a cube of piriform cortex tissue. Second, the number of neurons is reduced from the mouse to cat, leading to an increase in Glia to Neuron (GNR) ratios with an increase in brain volume. Notably, the number of glia and neurons change with brain regions: compare the piriform cortex (green) and neocortex (red) in mice. Glia densities can thus serve as region specific markers. Note that for ease of illustration the coronal section schematics for mouse and cat are not drawn to scale. The cat is drawn at half the actual scale, which means the cat should be twice as large as the schematic in the figure.

Interestingly, although more numerous, the volume density of glia in the human APCx was 42,000 glia/mm^3^, which is consistent with volume density of glia in other species (43,000 glia/mm^3^). Figure 6f shows that the human APCx glia density (in red) falls on the same regression line as other species (replotted Fig. 3c to include humans). This is because although the surface area density of glia doubles in humans compared to mice and other species, the corresponding thickness of the human PCx also doubles, leading to a similar volume density. Thus, although humans have the same volume density of glia, there are more glia in the human PCx because a larger piriform (by volume) will have a higher number of glial cells.

Mirroring the observed similarities in glia volume densities in APCx of mice and humans were glial density differences in between layers. In both, mice and humans (Figs. 2, 6), Layer 1 had significantly more glia than the other layers, and the cell-dense layer 2 had the fewest glia. An in-depth analysis of individual glial cell types shows that astrocytes, microglia, and oligodendrocytes also differ in their distributions between layers. In both species, most of the astrocytes were in Layer 1 (57% in mice, 41% in humans) followed by Layer 3 (26% in mice, 35% in humans), and Layer 2 (17% in mice, 24% in humans). Most oligodendrocytes, in contrast, had similar numbers in Layers 1 (39**%**, 50**%**) and 3 (43**%**, 27**%**), and fewer in Layer 3 (17**%**, 20**%**). Microglia were slightly different with both Layers 2 and 3 containing around 30%, with the rest (40 %) in Layer 1. Thus, glia as well individual glial cell type (spatial) distributions were broadly similar in mice and humans, suggesting glia as a whole might share some similarities in function across species.

To test if the spatial distribution patterns of glia from mice and humans are recapitulated in other species, we examined glia in each of the APCx layers. Just like in mice and humans, the highest number of glia were found in Layer 1 and the lowest in Layer 2 (Fig. S5c). We further examined the dataset to test if glial volume densities were different between the layers. As shown in Figure S5d, glia volume densities are highest in Layer 1 followed by Layer 3 and Layer 2, and the difference between Layers 1 and 2 is significant. Thus, glial distributions across layers might be preserved across species, from mice to humans. Notably, the distributions of neurons and synapses, too, vary, across layers (Neville and Haberly, 2004; Srinivasan and Stevens, 2018b), suggesting that differences in glia spatial distributions maybe a reflection of the different computations being carried out in each layer (Haberly, 2001b; Bekkers and Suzuki, 2013b).

## Discussion

We have identified two principles underlying glia organization in the brains of mammalian species (Figures 3 -- 5), which include rodents, carnivores, and primates including humans. First, glia volume density was constant within each brain region across brain sizes. This property was observed in four brain regions (Fig. 3c, anterior piriform cortex; Fig S3c, posterior piriform cortex; Fig. 4c, entorhinal cortex; (Carlo and Stevens, 2013), neocortex; Fig. S4b, cerebellum; Fig. S4d, comparison of all regions) that differ from each other in terms of morphology, connectivity, and function. Second, in the piriform cortex, neocortex, and the cerebellum, the number of glia per neuron increased with brain volume according to a ¼ power law (Figs. 5, S4, and equation 1).

### Consistency between individual and overall counts of glia types

In this study, we have used three methods to identify glial cells in the brain: DAB stains, Nissl stains, and sox9-eGFP mice. The first method that we used was DAB staining to identify three major glial cell types – astrocytes, microglia, and oligodendrocytes – and their densities in the piriform cortex of humans and mice. We, however, did not have access to corresponding tissue in other animals and to other regions in mice and humans, and, therefore, used Nissl stains. To test the effectiveness of using Nissl stains, we compared counts obtained by Nissl staining with those obtained by DAB staining (Figs. 2, 6 and Table 3). Estimates from Nissl stains were consistent with those obtained by DAB stains, in humans and mice, which gave us confidence to employ the Nissl stains to make cross-species comparisons. Notably, as the main goal of our study is the comparison between glia organization across regions and species, our results are valid, even if the absolute numbers with DAB stains were slightly lower (though, not statistically significant) than those reported with Nissl stains. It is perhaps one of the reasons that previous quantitative studies used similar methodologies (Carlo and Stevens; Friede, 1963; Sun et al., 2017) for cross-species comparisons, which would not have been possible otherwise.

**Table 3:**
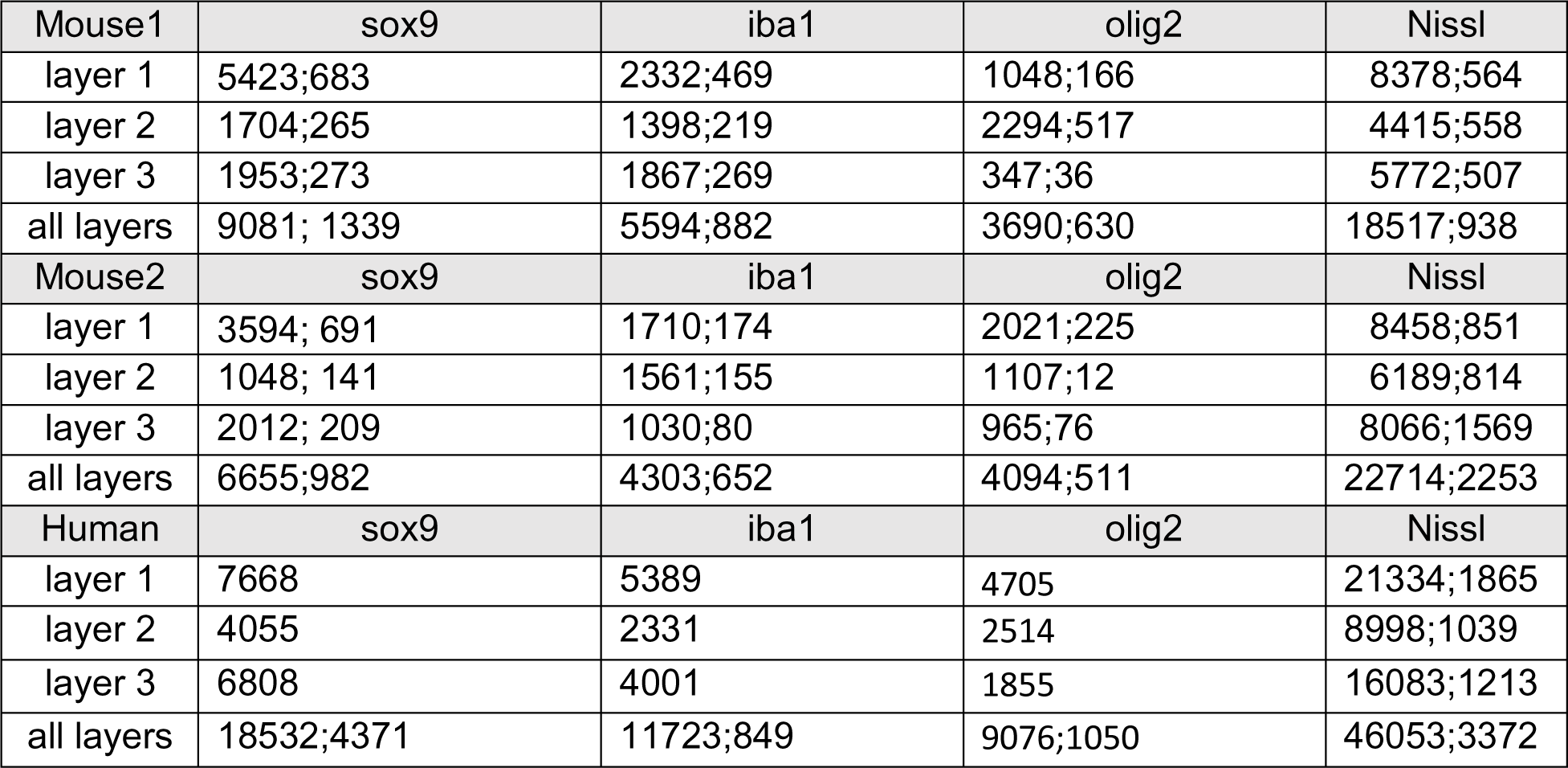
Summary of measurements in mice and humans with DAB and Nissl staining. For Nissl stains, surface area density numbers are giving as Mean, SEM of individual measurements. The SEM here is of the individual measurements for each stain type in each mouse.

The third method that we used – estimating astrocyte/neuron ratios through the expression of sox9 (to mark astrocytes) and NeuN (neurons) in SOX9-eGFP mice – provides additional support. The astrocyte/neuron ratios with SOX9-eGFP were consistent with those with DAB stains (Fig. S1). Finally, as elaborated in a later section, the proportion (astrocytes: microglia: oligodendrocytes) of individual glial cell types was similar across humans and mice providing additional validation for our DAB staining. Thus, the similarity between DAB and Nissl staining, and astrocyte/neuron ratios in SOX9-eGFP and DAB-stained mice provides strong support for using Nissl stains to estimated glial cell distributions in different brain regions across myriad species.

Notably, our investigation of individual glia types with DAB staining in mice and humans showed that, cumulatively, their count was slightly lower than estimates of overall counts obtained with Nissl stains. The cumulative count of the individual glial cell types accounted for about 81 – 85 % of the Nissl counts, respectively, in mice and humans. There are two reasons for the lower count. The first is that we used Olig2 to mark oligodendrocytes (Valério-Gomes et al., 2018). As previously noted, while an effective marker, this does not capture the entire population of oligodendrocytes, and there are few markers that do (Huang et al., 2022). Second, although not as numerous, ependymal cells, NG2, and radial glial cells, were not individually estimated but would have been part of the estimate through Nissl stains. Thus, the lower cumulative estimate with DAB stains is consistent with the counts obtained through Nissl stains.

It is important to note that besides glia demarcation into constituent cell-types, there is further sub-division possible as shown in numerous studies, for instance, astrocytes can be subdivided into at least interlaminar and protoplasmic astrocytes, which are found in distinct spatial contexts and likely serve different functions (Oberheim et al., 2009). Single cell studies have also exposed a diversity of astrocyte and other glial cell sub-types (Batiuk et al., 2020; Fang et al., 2022). Our results revealing a global property of glial cells is similar to studies of neuronal components in circuits, where workers have focused on measurements volumes and neuronal populations (Stephan et al., 1981; Finlay and Darlington, 1995b; Stevens, 2001) to enhance our understanding of neural circuits (Finlay and Darlington, 1995b; Zhang and Sejnowski, 2000; Striedter, 2004; Stevens, 2009). Our description of glial cells is a similar foray into the organization of glial cells, and emerging studies, e.g., single cell studies (Batiuk et al., 2020; Fang et al., 2022), can be combined with our work to enable finer-grained models of glial computations in circuits.

### Brain areas can be distinguished based on their glial density

As noted above, although constant within a brain region, glia volume densities differ between regions (Fig. S4d). The neocortex and APCx illustrate the differences well: the volume occupied by a glial cell in APCx is smaller. Indeed, each glial cell in APCx occupies a neuropil volume covering 0.022 nL, which corresponds to a 28 µm cube on each side. This is significantly smaller than the volume occupied by a glial cell in the neocortex (Carlo and Stevens, 2013), where, on average, the territory covered by each glial cell is a cube with side 36-40 µm (Ogata and Kosaka, 2002; Halassa et al., 2007; Oberheim et al., 2012). Glia density in PPCx is midway between the two regions, with glia occupying 0.03 nl of volume, which corresponds to a cube 32 µm on each side. In addition, in ECx, which is a six-layered structure, each glial cell covers a territory equivalent to a cube that is 46 µm on each side, closer to the six-layered neocortex.

It is plausible that the constancy in glia density is not exclusive to the cerebrum. This possibility is supported by our re-analysis of a study by Friede (Friede, 1963), who examined the cerebellum of 18 species ranging from birds and rodents to bigger mammals, and estimated the number of glial cells per mm^3^ of the molecular layer of the cerebellum, as well as the number of Purkinje cells. Although, in his interpretation of the data, the author concluded that glia densities were different between species, when we replotted the data from the study (leaving out five species of birds (non-mammals) and rodents in which glia densities were intriguingly higher), we found that glia densities were, in fact, normally distributed around a mean of 22,840 glia/mm^3^ (Fig. S4 a, b, c). This value, with an associated SEM of 1,592 likely represents the true density of glia in 13 of the 18 species studied, all of them mammals (Fig. S4a, b, c). This glial volume density corresponds to glia occupying a cube of side 36 µm. Of course, we do not suggest that all glia are not uniformly spread apart, but, rather suggest that individual glial-cell types or even glial-cell subtypes might spread themselves evenly to uniformly sample space as studies have suggested (Bushong et al., 2002). If the proportion of glial cell-types are maintained across species (as our APCx data suggests in Figs. 2, 6), then the cube sides that we showed above would be changed by a constant factor, while maintaining relative differences (would have the same ratio) across individual regions.

### Glia densities as markers of circuit differences

What are the factors determining glial densities within regions? Although, outside this study’s scope, prior work suggests that each region’s unique glial density might arise due to differing functional demands. There are two examples of function determining glial density. The first example comes from aging studies in mice, rats, and humans. In humans, aging was shown to promote a marked increase in the number of microglia in regions associated with immune responses and inflammation (Soreq et al., 2017). Similarly, studies done in rats show that, with aging, the number of microglia increased (about 16%) in the frontal and parietal regions without a significant change in the numbers of neurons, although axon and dendrite degradation was observed (Peinado et al., 1993, 1997). As glial cell types come in different sizes, with astrocytes, in general, being considerably larger than most other types of glia (Rajkowska et al., 1998; Oberheim et al., 2012; Davis et al., 2017), a change in glial cell type composition could change glial density.

A second example of glial densities reflecting differing demands is the greater density of glia in both APCx and PPCx compared to the neocortex. The organization of the connection matrix from the olfactory bulb to PCx in the mouse shows that every glomerulus contacts every piriform neuron (Srinivasan and Stevens, 2019). This all-to-all connectivity calls for a large number of synapses, possibly more than a similarly sized circuit in the neocortex, where circuits are topographic and the number of synapses required might be fewer, or, at least, distributed over a larger cortical thickness with its six layers. Studies have shown that the molecular layer of the piriform cortex (layer 1) is particularly synapse-rich (Schikorski and Stevens, 1999; Neville and Haberly, 2004; Srinivasan and Stevens, 2018b). Measurements of glial surface density (Table S1) support these studies: across species, layer 1 of the piriform cortex has a greater number of glial cells (Figs. 2, 5, S5). Thus, it is likely that the synapse-dense layers of the piriform cortex might require support from a higher number of glia, in agreement with observations (Faraguna et al., 2010).

Work in the last decade showing abrupt astrocyte boundaries between regions (Eilam et al., 2016) supports our thesis that glia densities are similar within a region across species, but differ from one region to another. Eilam et al. showed that astrocyte processes seem to taper off at the boundaries of barrel cortex, auditory cortex, and visual cortex. Related studies have found that astrocytes play an important role in integration of local neuronal processing (Araque et al., 1999; Haydon and Carmignoto, 2006). Astrocytes collect information about processing in their local vicinity by sampling synapses, integrate this information, and channel it to blood vessels (Attwell et al., 2010; Eilam et al., 2016). Taken together these studies provide evidence that the makeup or spatial distributions of astrocytes reflect local computations within a circuit. Given that neuronal makeup and processing varies with each circuit, their astrocyte makeup would vary, too. If other glia have similar physical constraints, it might suggest that glia within a region exclusively serve a local regional function, and thus have differing densities.

### Glial Scaling

The conservation of glial density provides yet more evidence that circuits are optimally designed. Extra neuronal machinery increases energy usage at the cost of diminishing returns and is not evolutionarily advantageous. A similar explanation could be extended to glial organization. As we know, various glial sub-classes handle important tasks such as maintaining efficient synaptic plasticity and transmission, regulating blood flow, and clearing waste products. Each region’s neurons are distinctly organized (in their sub-types and connection structure) to fulfill specialized computational tasks. Correspondingly, it is possible that glia, which enable these computations, are optimally balanced, and any additional glial machinery will add to the system’s cost while providing diminishing benefits. Thus, glia organization is likely to be precisely engineered (reflected in conserved glial constraints) for the optimal function of neural circuits, just like neuronal organization (Sterling and Laughlin, 2015).

It is worth noting that the scaling relationships examined in our current work included primates and non-primates. For instance, for APCx, PPCx, and frontal cortex (Sherwood), primate brains follow the same constraints on glia organization as other animals. Even if there are species-specific differences at the molecular or cellular level, at the population level, glial organization is similar. Thus, population-level findings in model animals like mice could apply to primates as well and make these studies more relevant for human neurophysiology.

While we have unearthed broad principles of glia conservation in terms of their spatial distributions, their number with respect to neurons, and distributions of individual glial cell types, there are documented morphological differences in glial cells across species, too, and is reflected in their function. An in-depth study of astrocytes in mice and humans has shown that human astrocytes are larger and have more numerous processes (Oberheim et al., 2009). These differences might be one part of the reason why their functions might differ, for instance, in human astrocytes, calcium propagation is faster (Oberheim et al., 2009). And, human astrocytes lead to better function when transplanted in mice (Han et al., 2013). Additionally, single cell studies have highlighted that within the same region, but in different species, the sub-types of each glial cell type could also differ mirroring earlier studies showing that primates have two types of astrocytes -- polarized and interlaminar -- not observed in rodents (Oberheim et al., 2009). Similar findings in ferrets (Lopez-Hidalgo et al., 2016) suggest a possible trend across species. Lastly, the glia covered in this study are part of the grey matter. It is possible that the spatial distributions of white matter glia will diverge from grey matter glia and they might even differ across species reflecting differences in connectivity between species. An experimental approach to the question of how glia may contribute to high cognitive abilities is to follow up on studies that have grafted human glial cells into mouse cortex and reported faster learning and enhanced long-term potentiation (Han et al., 2013). Additional testing might reveal further differences.

### Glia-neuron ratios increase with brain volume according to a ¼ power law

Our findings show that glia, like neurons, are constrained by orderly relationships within circuits. This constraint reinforces the idea that there is a strong functional coupling between glia and neurons, which is integral to the functioning of the circuit. By analyzing GNRs, we aimed at gaining new insight into those interactions. Our determination of GNRs also provides important constraints for realistic *in silico* models of glia-neuron interactions.

GNR increased with brain volumes according to a ¼ power-law in four brain regions. These include the anterior piriform cortex (from measurements in this study and (Srinivasan and Stevens, 2019)), the prefrontal cortex of primates (analysis of data from (Sherwood et al., 2006)), cerebellum (analysis of data from (Friede, 1963)), the cerebral cortex as a whole (Haug, 1987; Verkhratsky and Butt, 2018), and four regions of the neocortex (not including prefrontal or visual cortices) in mammals based on an investigation of glia and neuronal constraints in the neocortex uncovered by (Carlo and Stevens, 2013).

One question arising from these observations is how bigger brains came to have more glia per neuron. As brains grow bigger, there are several changes in a neuronal network with respect to numbers of neurons, the connections they make, and the synapses borne out of these connections. First, in bigger brains, neurons have to communicate over wider distances, both locally, and long-range to transmit information in a comparable amount of time to smaller brains (Buzsáki et al., 2013). As a result, the width or diameter of their axons increases to accommodate greater conduction velocities (Hursh, 1939; Ritchie, 1982; Sakai and Woody, 1988; Innocenti, 2011), leading to a greater amount of wire in extracellular space. Bigger axons also necessitate bigger soma (Sakai and Woody, 1988; Herculano-Houzel, 2014). These factors increase glia sizes as well (though not to the same extent). All these changes result in an overall increase in volume. Previous studies have shown that the number of neurons under a square mm is constant across species (Rockel et al., 1980; Carlo and Stevens, 2013). In order to accommodate the same number of neurons under a square mm, the thickness of the cortex would need to expand to increase volume as observed (Stolzenburg et al., 1989; Carlo and Stevens, 2013). The result is that glia volume density stays constant while neuronal volume density decreases, leading to higher GNRs with bigger brains.

### Consequences for glia tiling across species

In this paper, we have shown how glial densities are conserved within a circuit across species. Indeed, in mice and humans, we showed that astrocyte densities are similar, too. Studies of astrocytes tiling a region and apportioning equal amounts of territories in the mouse hippocampus support this view (Bushong et al., 2002; Halassa et al., 2007). Bushong et al. also showed that in mice, the amount of overlap between neighboring tiled regions is minimal. These findings speak to existing theories of how astrocytes support computations within their immediate neighborhood. Recent studies, however, have shown that the volume of space occupied by astrocyte soma as well as their processes increase with bigger species. In mice, ferrets, humans, and non-human primates (macaques and chimp), astrocyte size (soma as well as overall volume including those of processes) increases with species brain volume (Oberheim et al., 2009; Verkhratsky and Butt, 2013; Lopez-Hidalgo et al., 2016). As astrocyte densities are conserved across species (Figs. 2, 6), their soma must be regularly distributed in space, suggesting that as their volume becomes larger, the area of overlap in their territories must increase. Recent work comparing larger brained species to mice showing a greater overlap between neighboring astrocytic processes (Oberheim et al., 2009) are consistent with implications of conserved density that our study revealed.

### Glia and cognition

It has been hypothesized that the increase in GNRs observed in species with bigger brains positively correlates with cognitive abilities, which even led to similar claims for humans. For instance, it was reported that Brodmann area 39 of Einstein’s brain, part of the parietal lobe, housed a higher number of glia per neuron than eleven other male subjects (Diamond et al., 1985). Evidence in support of such an observation, however, relied on an *n* of 1 (Einstein’s brain). Moreover, while GNRs in Brodmann area 39 were higher for Einstein, they were just one of many factors studied, all others not being significantly different from the control population. When comparing multiple variables, it is possible that one of the variables is statistically significant by chance. To establish or reject the notion that higher GNRs lead to greater cognitive abilities, one would need to systematically analyze the possible correlation between cognitive abilities and GNR, e.g., investigate whether individuals with high cognitive abilities have above average GNRs and vice-versa. These studies could then lead to the question of whether both characteristics arise from a common root and/or are causally related.

### Conclusions

Previous studies of neuronal scaling (Finlay and Darlington, 1995a; Zhang and Sejnowski, 2000; Sherwood et al., 2006; Stevens, 2009; Herculano-Houzel, 2010; Srinivasan and Stevens, 2019) which linked structure and function, did so by considering neurons as a single class. It was a necessary first step, as it helped identify general principles of neural function (Finlay and Darlington, 1995a; Stevens, 2009; Sterling and Laughlin, 2015), and paved the way for more complex models of how individual cell types within circuits contribute to function (Hodge et al., 2019; Bakken et al., 2021). Similarly, to derive high level insights into the role of glia in neural function, we lumped glial cell-types together, similar to previous studies (Stolzenburg et al., 1989; Sherwood et al., 2006; Carlo and Stevens, 2013). Our results showing that glia organization in a brain region is conserved across different brain sizes sets the stage for further in-depth studies of roles of specific glia types as well as constraints on them. Building on this finding, we investigated the composition of individual glial types in the piriform cortex of humans and a model organism central to neuroscience – the mouse – to show that while glia compositions are similar, total glial numbers differ between the two species. An intriguing possibility warranting future investigations is that different glia compositions in topographic (e.g., neocortex), distributed circuits (e.g., piriform cortex), and circuits with a mixed architecture (e.g. entorhinal cortex) might account for differences in glia density and in their specific functions.

## Acknowledgements

We thank Scott Magness for providing us with Sox9-EGFP mice, Nicola Allen, Ashley Brandebura, Phong Ngyuen for technical and reagent help for the DAB experiments, and Jorge Aldana for computational assistance. We also thank Chuck Stevens, Nicolla Allen, and Sinda Fekir for valuable feedback on the study. This research was funded in part by the Howard Hughes Medical Institute, Kavli Institute for Brain and Mind, and NIH DC017695.

