## Supplementary Information for "Laws for Glia Organization Conserved Across Mammals"

### Supplementary Materials

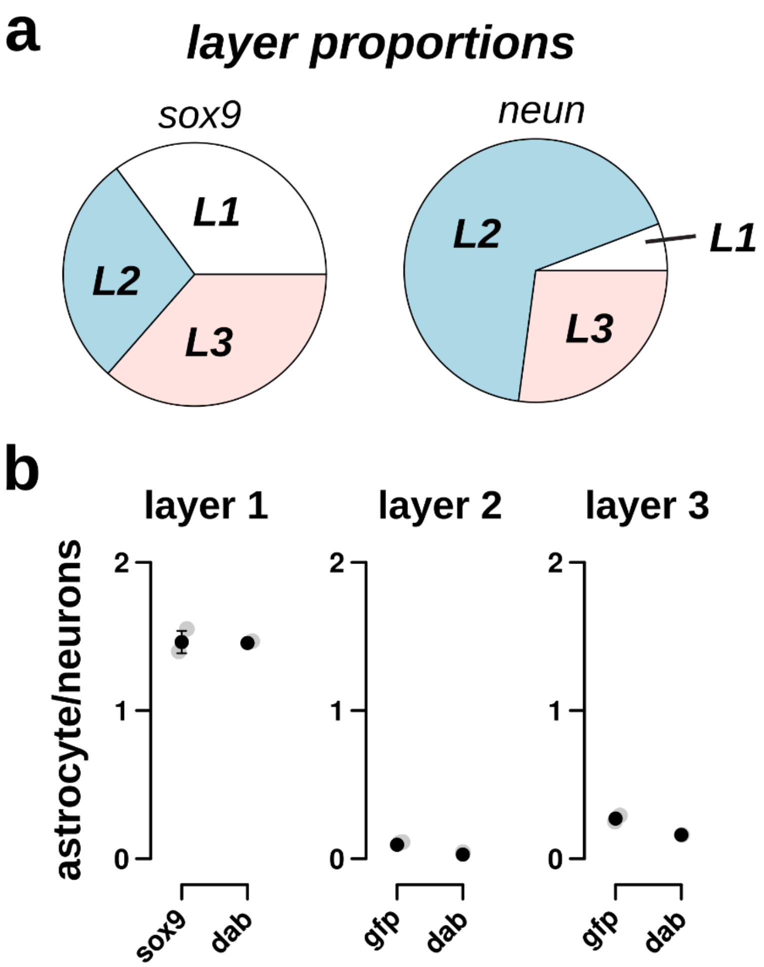


**Figure S1: DAB staining estimates of astrocytes are consistent with sox9-EGFP mice estimates.** (a, left) Pie-chart displaying the proportion of astroglia (with sox9-GFP) in the three layers of the piriform cortex. Layer 1 has more astroglia than layers 2 and 3. (a, right) Proportion of neurons labeled with NeuN across layers in Sox9-GFP mice. (b) Astrocyte-to-neuron ratios with sox9-GFP and sox9-DAB staining across the three cortical layers. The highest ratio were observed in layer 1, followed by layer 3 and, finally, layer 2. Layer 1 has fewer neurons, containing more astrocytes. Error bars are confidence intervals. Data for neurons in this plot is from (Srinivasan and Stevens, 2018).

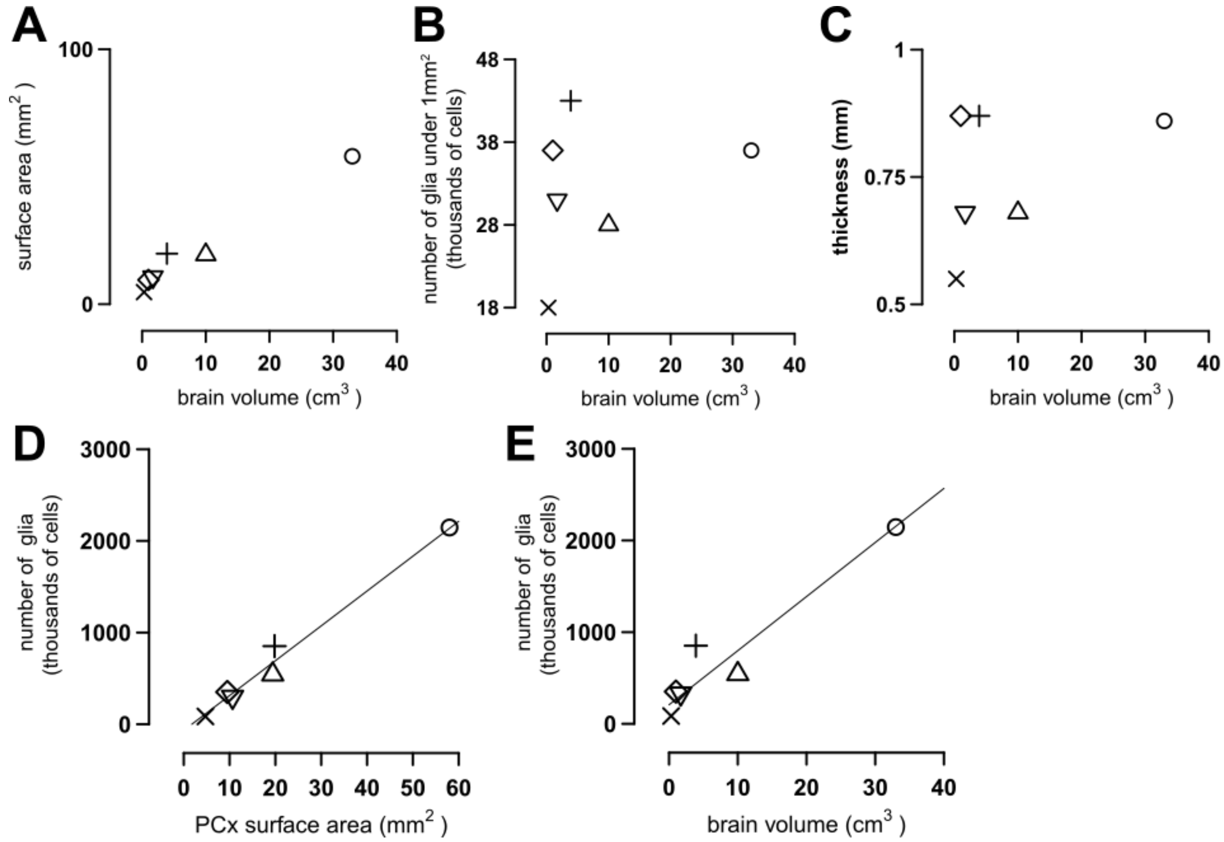


**Figure S2: Summary of APCx measurements across species.** The number of glial cells was plotted for six species of mammals: mouse (×), rat (▽), ferret (D), opossum (◊), cat (o), and guinea pig (+). (A) The surface area of the APCx increases with an increase in brain volume (cm$^3$). (B) The number of glia underneath a square mm of APCx versus brain volume. (C) The thickness of the APCx versus brain volume. (D) Absolute number of glial cells plotted against the size (surface area) of the APCx ($R^2$ = 0.98, Pearson correlation coefficient). The regression line is described by the equation: y = -67 + 38x, where y is the number of glia (in thousands) and x is the APCx surface area in mm$^2$. (E) Absolute number of glial cells in the APCx plotted against brain volume ($R^2$ = 0.86, Pearson correlation coefficient). The regression line is described by the equation: y = -242 + 59*x, where y is the number of glia (in thousands) and x is the brain volume in cm$^3$.

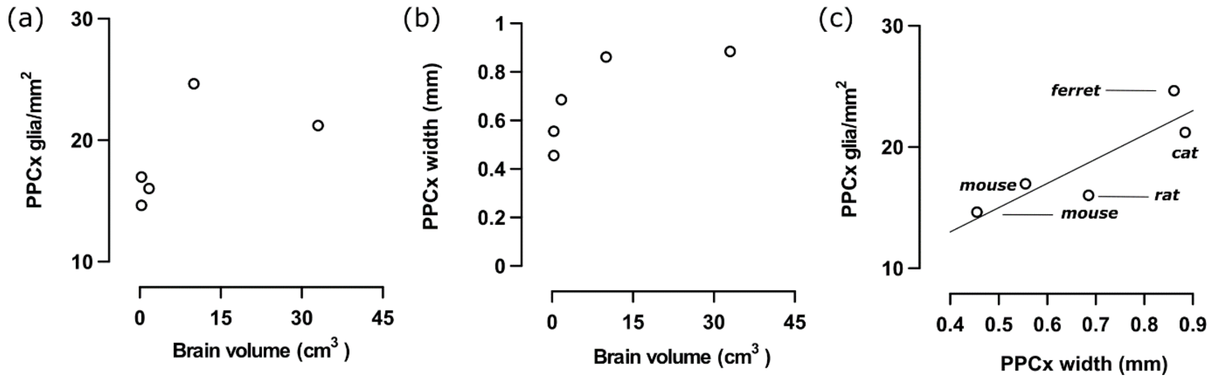


**Figure S3: Glia volume densities are similar in PPCx across species.** (a) As the brain volume of the four species studied (mouse, rat, ferret, cat) increases, so does the number of glia under 1mm$^2$ of PPCx surface. (b) Similarly, as brain volume increases, so does the width of the PPCx. (c) With an increase in the width of the PPCx, the number of glia also increases, leading to similar volume densities across species. The fitted line has the form $y = 20x + 5$, and an $R^2$ of 0.76. The volume density shown in Fig.

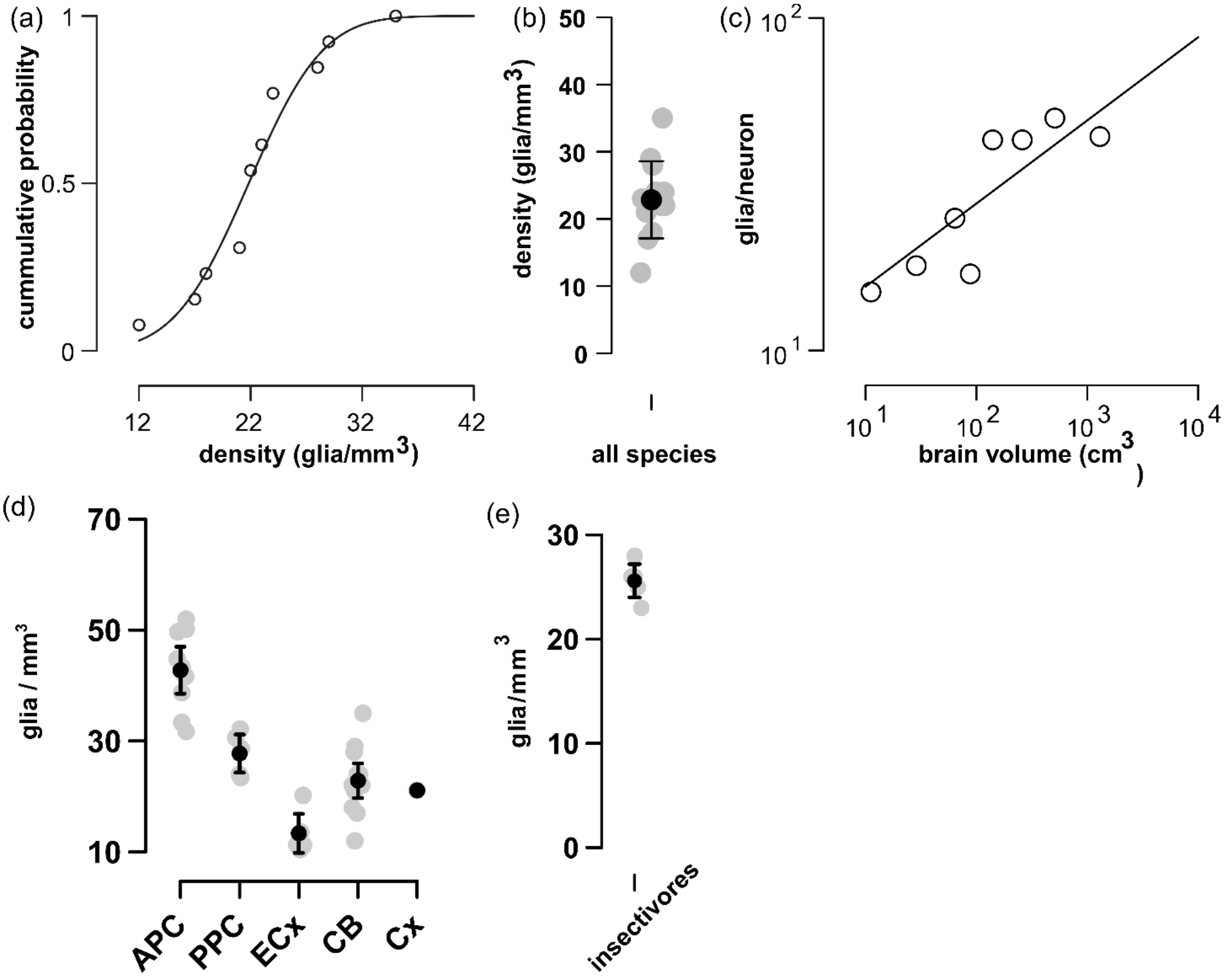


**Figure S4: Glia volume density and glia-neuron ratios in the cerebellum of multiple mammalian species.** The glia volume densities (i.e. number of glia per unit volume) in twelve species (human, monkey, cow, horse, sheep, elk, pig, dog, deer, lion, cat, rabbit, and frog) as reported by (Friede, 1963) were replotted as a cumulative probability curve (a). Note that the distribution of glia density follows a Gaussian distribution tightly centered around a mean value of 22.8 glia/mm$^3$ with an SD of 5.74 (a,b), suggesting that such a value represented, in fact, a conserved across the species studied. Panel (c) shows that, as brain size increase (plotted in the order of rabbit < cat < sheep < dog < lion < horse < man), cerebellar glia-neuron ratio increased according to the relationship, $y = 0.3\ x^{1/4}$ with an $R^2$ of 0.68 and CI of 0.14-0.29. Note that brain volumes in (c) were taken from (Haug, 1987), and, for the sake of consistency, we only plotted in this panel species that overlapped between that study and (Fried, 1963). (d) Comparison of glia volume densities in five regions: anterior piriform cortex (APC), posterior piriform cortex (PPC), entorhinal cortex (ECx), cerebellum (CB), and neocortex (Cx). The plot shows that the glia volume densities of each region are distinct; compare the confidence intervals which are non-overlapping between APC, PPC, and ECx showing evidence that the mean glial volume density for each region are significantly different. Note that volume densities in the neocortex and cerebellum seem similar, and that of the cerebellum overlaps with PPC.

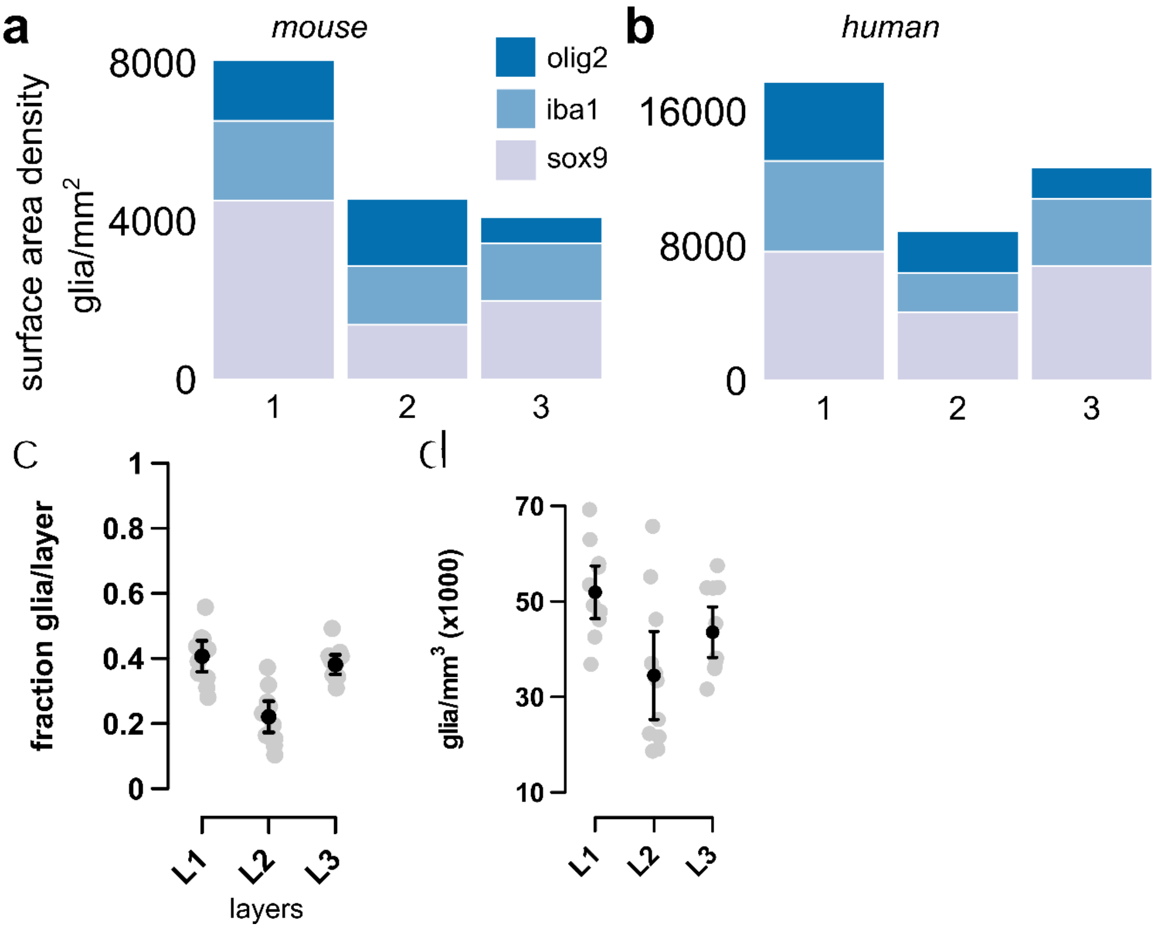


**Figure S5: Distribution of astrocytes, microglia, and oligodendrocytes across anterior piriform cortex layers.** (a) The distribution of the three major glial cell types in the mouse piriform cortex. Most (> 50 %) of Layer 1 glial cells are astrocytes, while about 50% of layer 3 cells are astrocytes. The three cell types are equally distributed in layer 2, which also contains the most neurons. (b) The distribution of the three glial cell types in the human piriform cortex. Astrocytes form the greatest proportion of cells in all three layers, though, unlike the mouse, their proportion is lower in layer 1, which contains the most glial cells of all layers. (c) Number of glia in each layer varies, with Layer 2 having the lowest and Layer 1 having the most. (d) The volume density of glia is highest in Layer 1 and lowest in layer 2 with the difference being significant. In both (c) and (d) Layer 3 has an intermediate number of glia and glia volume density.

*(d) The volume density within layers of APCx is similar, across species. Each point on the plot represents the width of the layer (1—3) versus the number of glia under a square mm within that layer. We plotted all layers for all species. The fitted line $y = 0.8 + 47x$ shows that the number of glia is proportional to the width of the layer ($R^2 = 0.6$). See supplementary Tables 1 – 3 for values of widths and densities.*

**Table S1.** The surface area density (the number of glia underneath a square mm) of the APCx. We also present results for individual layers. All values are given in terms on number of glia/mm$^2$.

| Species' common name | Total density ( glia/mm$^2$ ) | Layer 1 | Layer 2 | Layer 3 |
|---|---|---|---|---|
| **Cat** | 37507 | 17280 | 4977 | 15249 |
| **Ferret 1** | 27606 | 11821 | 4226 | 11559 |
| **Ferret 2** | 36708 | 10293 | 13671 | 14158 |
| **Guinea pig 1** | 43942 | 15581 | 11108 | 17251 |
| **Guinea pig 2** | 43522 | 14834 | 13870 | 14818 |
| **Mouse 1** | 18517 | 8378 | 4415 | 5722 |
| **Mouse 2** | 20393 | 7968 | 5414 | 7010 |
| **Opossum** | 37427 | 11610 | 8660 | 18424 |
| **Rat 1** | 31788 | 17722 | 3275 | 10998 |
| **Rat 2** | 24005 | 12963 | 3250 | 7963 |

**Table S2.** Summary of APCx measurements across individual layers. All values are in mm.

| Species' common name | Total width (mm) | Layer 1 | Layer 2 | Layer 3 |
|---|---|---|---|---|
| **Cat** | 0.8793 | 0.2744 | 0.1971 | 0.4077 |
| **Ferret 1** | 0.6630 | 0.2248 | 0.2225 | 0.2194 |
| **Ferret 2** | 0.70666 | 0.2422 | 0.2082 | 0.2561 |
| **Guinea pig 1** | 0.8760 | 0.2723 | 0.3023 | 0.3012 |
| **Guinea pig 2** | 0.8151 | 0.2500 | 0.2924 | 0.2727 |
| **Mouse 1** | 0.5553 | 0.1761 | 0.1988 | 0.1813 |
| **Mouse 2** | 0.5266 | 0.1683 | 0.1656 | 0.1968 |
| **Opossum** | 0.8639 | 0.2545 | 0.1605 | 0.4838 |
| **Rat 1** | 0.7194 | 0.2482 | 0.1490 | 0.3121 |
| **Rat 2** | 0.6483 | 0.2453 | 0.1805 | 0.2252 |

**Table S3.** Estimates of number of glial cells per unit volume (volume density) in the APCx based on direct measurements.

| Species' common name | Vol. density (glial cells/mm$^3$) |
|---|---|
| **Cat** | 42,660 |
| **Ferret 1** | 41,598 |
| **Ferret 2** | 52,066 |
| **Guinea Pig 1** | 49,800 |
| **Guinea Pig 2** | 56,000 |
| **Mouse 1** | 34,020 |
| **Mouse 2** | 40,693 |
| **Opossum** | 43,800 |
| **Rat 1** | 45,000 |
| **Rat 2** | 36,861 |
| **Mean ± SEM** | 44,249 ± 2,140 |

**Table S4.** Number of cortical columns analyzed per animal for APCx.

| Common name | # of columns |
|---|---|
| Cat | 10 |
| Ferret 1 | 16 |
| Ferret 2 | 10 |
| Guinea Pig 1 | 11 |
| Guinea Pig 2 | 9 |
| Mouse 1 | 24 |
| Mouse 2 | 8 |
| Opossum | 16 |
| Rat 1 | 15 |
| Rat 2 | 21 |
| **Mean ± SEM** | 14 ± 1.7 |
